# Clinical breakthrough: Biomimetic Scaffolds Based on Soluble Collagen for Regenerating Large Volumes of Highly Functional Tissues

**DOI:** 10.1101/2025.06.19.660623

**Authors:** Xiaoming He, Zhaohui Luo, Shenghua He

## Abstract

Functional tissue regeneration in tissue engineering has long been an aspiration of humanity. However, conventional materials such as metal, ceramic, polymer, and decellularized extracellular matrix (dECM) have not yet yielded the desired outcomes.

We have developed a collagen biomimetic material based on soluble collagen. This biomimetic material emulates the composition, structure, and function of native tissue. It possesses high strength, high affinity, high nutrient, and low antigenicity. The biomimetic material had been utilized to fabricate some implants (absorbable surgical suture, artificial tendon, hernia patch) and had demonstrated remarkable functional regeneration in animal experiments. In our previous researches, collagen surgical sutures facilitated wound healing, artificial tendons and artificial muscles (hernia patches) successfully repaired extensive defects of tendons and muscles, respectively. These successful tissue engineering endeavors enabled damaged defect to restore its biological functions and structure. Degradation of these biomimetic implants synchronized with the regeneration of the new tissues.

Based on these successful functional regenerations, the present study focuses on regeneration of rabbit sciatic nerve. This collagen biomimetic material was employed to construct an artificial nerve tube (length: 2 cm, inner diameter: 2 mm, outer diameter: 3.5 mm) to repair a 2 cm-long defect in the rabbit sciatic nerve. After 36 weeks, the newly formed functional sciatic nerve successfully bridged the damaged ends, while approximately 92% of the artificial nerve tube has degraded.

In this study, we provide an overview of the biomimetic material and its medical applications. We believe that these successful regenerations of tendon, muscle, and nerve all stem from the exceptional characteristics of this biomimetic material. We firmly believe that this collagen biomimetic material exhibits remarkable regenerative capabilities in diverse tissues. It can pave the way for the fabrication of increasingly critical artificial implants, which can be utilized to repair defects in human tissues, including artificial esophagi, tracheae, and small blood vessels. In regenerative medicine of induced pluripotent stem cells (iPS), the biomimetic materials can replace temperature-sensitive materials (PIPAAm) to fabricate cell sheets. This approach expedites the process, enhances cellular vitality, and consequently improves regeneration outcomes.

This material technology not only provides a revolutionary solution for clinical treatment but also paves the way for a new development direction in regenerative medicine.

## Introduction

For irreversible tissue damage or deletion resulting from diseases, accidents, or hereditary factors, autologous tissue collection and transplantation remain the gold standard. Autologous tissue exhibits exceptional biological compatibility and regenerative potential. However, the availability of autologous tissue is frequently inadequate[Langer and Vacanti, 1993; Atala 2004; Vacanti and Bonassar 2000; Griffith and Naughton, 2002].

Currently, commonly utilized alternative materials in tissue engineering include metals, ceramics, synthetic polymers, and dECM. However, these materials exhibit significant compositional, structural and functional differences from those of native tissues [Langer and Vacanti, 1993; Place et al.,2009; Chen and Liu, 2016]. For example, metals and ceramics have favorable mechanical properties, simultaneously they exhibit pronounced biological inertness, limited affinity, and susceptibility to thrombosis, inflammation, and local tissue fibrosis [Chen and Liu, 2016].

Natural polymers such as collagen, gelatin, demonstrate favorable biocompatibility; however, their degradation is rapid or lacks adequate mechanical strength. In contrast, durable synthetic polymers can generate detrimental degradation by-products or induce chronic inflammation, exemplified by polylactic acid (PLA) and polycapenal ester (PCL)[Langer and Vacanti, 1993; Baker and Mauck, 2007; Li et al., 2002]; dECM of animal tissues, including pig pericardium and cow tendon, are employed for valve or soft tissue repair after reducing immunogenicity by decellurization. Nevertheless, non-helical telopeptides and residual antigen components (e.g., DNA, heterogens) may still elicit immune responses, chemical residues may also induce inflammatory response [Crapo et al., 2011; Keane and Badylak, 2015]

Furthermore, over time these implants in vivo are frequently loosened, worn, or corroded due to humor erosion or mechanical friction, which escalates postoperative complications and necessitates secondary surgery to remove the defective material [Williams, 2008].

Consequently, the clinical medical community urgently seeks to develop regenerative medical materials with high affinity, low antigenicity, exceptional biodegradability, and high strength that akin to autologous tissue, can replace conventional materials. However, this objective has not yet been achieved to meet the pressing clinical requirements.

Except the material component, the remarkable complexity of hierarchical structure of native tissue is perhaps also another important factor for successfully functional regeneration. For instance, tendons and ligaments are composed of highly regularly aligned fibers, facilitating stretching. Furthermore, the regeneration and performance of functional tissues necessitate extensive vascularization and innervation [Rouwkema et al., 2008; Debbi et al., 2022]. Consequently, structural refinement techniques such as 3D printing and electrospinning technology are utilized to mimic the natural arrangement of collagen fibers [Li et al., 2002]. Additionally, stem cell engineering, particularly mesenchymal stem cells (MSCs) were untilitazed to improve regeneration, and the implantation of bioactive molecules like IGF-1 and VEGF into scaffolds are employed to promote angiogenesis and induce regeneration [Rouwkema et al., 2008; Chen and Liu, 2016; Park et al., 2015]. Despite these efforts, substantial progress has not been made, particularly in the realm of functional loading tissues such as muscles, tendons, and ligaments. These tissues persistently present challenges in the successful regeneration process following significant deletions.

Collagen, a fundamental structural protein in the human body, plays a crucial role in numerous vital metabolic processes. Soluble collagen derived from animals serves as an exceptional medical material due to its low immunogenicity and high affinity. However, its mechanical strength is significantly compromised, and it undergoes rapid degradation in vivo. Researchers have employed various techniques, such as enzyme cross-linking, light cross-linking, glutaraldehyde treatments, and compounding with elastin, silk protein, and other materials, to enhance the mechanical properties and stability of collagen scaffolds [Lee et al., 2011], however, significant progress has not been realized.

Based on these previous research, we have developed an innovative medical material derived from soluble collagen. This medical material closely resembles the composition, structure, and functions of biological tissue. It exhibits exceptional strength, low antigenicity, and high affinity, making it particularly valuable for its regenerative potential in repairing tissue defects of the human body.

The biomimetic medical material, employed as absorbable surgical sutures, has demonstrated efficacy in facilitating healing skin wounds in both rabbits and humans [He et al., 2025a]. Additionally, it has exhibited successful functional regeneration of volumetric muscle loss (VML) and has successfully regenerated 2 cm Achilles tendon defects in rabbit models [He et al., 2025b; He et al., 2025c]. Notably, nerve regeneration was also successfully achieved in this current study, underscoring the material’s comprehensive application potential in the field of regenerative medicine.

This research provides an overview of innovative biomimetic materials and their applications as absorbable surgical sutures, artificial tendons, artificial muscles, and their current applications in nerve regeneration. It emphasizes the pivotal role of biomimetic materials in unlocking the regenerative potential of autologous stem cells.

Advancements in material technology will significantly enhance regenerative medicine, both in clinical practice and scientific research.

## Materials and Methods

### 1. Fabrication of artificial implants

#### 1.1. Fabrication of collagen threads (sutures)

Collagen threads are primarily fabricated from soluble collagen. Fabrication details are provided in [He et al., 2025a]. The properties of the collagen thread meet the standards of absorbable surgical sutures, successfully closing skin wounds in animal experiments and preliminary human applications.

Artificial tendons and hernia patches have been fabricated by weaving collagen threads. Notably, artificial tendons were successfully employed to repair a 2-cm tendon defect, as detailed in [He et al., 2025b]. Collagen hernia patches were successfully utilized to repair VML in the rabbit abdomen, as further elaborated upon in [He et al., 2025c]. The following are concise in terms of fabrication methods.

#### 1.2. Fabrication of artificial implants

##### 1.2.1. Artificial Tendons

(length: 2cm; diameter: 2mm): Collagen sheets of 12×20mm are fabricated by weaving collagen threads(diameter 0.080), then the sheet is rolled to be artificial tendons.

##### 1.2.2. Hernia repair patches

(30×30mm sheet, thick:0.4mm): the sheet is woven by collagen threads (diameter 0.15mm).

##### 1.2.3. Nerve guide tube

(length:2cm, inner diameter: 2mm; outer diameter:3.5mm): The tube is woven using collagen threads (diameter 0.10mm). The weaving method is analogous to that employed for collagen hernia patches.

### 2. Animal experiments

#### 2.1. Animals

NewZealand white rabbits weighing 2.0∼2.7kg were from Hubei Experimental Animal Research Center.

#### 2.2. Animal ethics

All animal procedures were conducted in accordance with applicable laws, regulations, and institutional guidelines for laboratory animal use and care. Animal welfare and ethical considerations were strictly adhered to. The laboratory animals were housed in a 12/12 light cycle environment with a temperature of 22±1°C and a humidity of 50%. Food and water were provided on an on-demand basis. The animal experiments were approved by the Ethics Committee of Wuhan University Central South Hospital (No. 2019007). Animals were euthanized by administering an overdose of anesthesia.

#### 2.3. Anesthesia and disinfectant

General anesthesia was administered during the surgical procedure. A 10% hydrated chloraldehyde peritoneal injection, administered at a rate of 1 mL per kilogram of body weight, was combined with a fast-sleep-new II intramuscular injection, administered at a rate of 0.3 mL per kilogram of body weight. The surgical area was thoroughly shaved and disinfected with iodine. Immediately following the operation, penicillin sodium (200,000 units per rabbit) was administered intramuscularly and continued for a duration of three days.

### 3. Animal surgery-treating different models 3.1-Skin wound model

Test suture (collagen suture) and control (PLGA) suture (w9915, Ethicon) were used to close 3-cm incisions on the rabbit’s back. Preliminary human application was also implemented: collagen surgical suture was used to suturing an incision wound after the removal of a prosthetic from a woman breast.

#### 3.2 Tendon defect model

Artificial tendons were used to repair 2cm long tendon defect.

#### 3.3 -VML model (hernia model)

Collagen hernia patch was used to repair 3×3cm VML hernia model,

#### 3.4. Sciatic nerve defect model

A linear longitudinal postero-lateral incision was made in lateral thigh by a #15 blade with the rabbits in right lateral recumbency. Then the fascia is cut open and smooth dissection performed between the anterior and posterior muscular compartments. Clear visual field of the sciatic nerve appears. A 2-cm nerve section was cut off with a scissor. Then an artificial nerve-tube (2mm inner diameter) was connected to the nerve defect. The nerve stumps were gently inserted inside the tube and fixed by using the 7-0 nylon suture. The fascia was closed by 5-0 nylon suture, and a skin closure is with 3-0 nylon suture.

Five rabbits underwent random euthanasia due to excessive anesthesia at 12, 24, and 36 weeks following the surgical procedure. The surgical site was incised to observe and document the surrounding tissue. This procedure assessed the healing process of the anastomosis and nerve regeneration. Subsequently, the transplanted nerve and newly formed tissue were meticulously excised for the determination of degradation rates, mechanical strengths, and the histological analysis. At 4 and 8 weeks post surgery, the surgical conditions were also observed by sacrificing 1-2 rabbits.

### 4. Measurement

#### 4.1. Tensile strength and Suture Strength

Strength of native sciatica nerve of rabbits and artificial nerve in vivo were measured, as detailed in [He et al., 2025b, 2025c] for measuring artificial tendon, hernia patch.

#### 4.2. Degradation

The degradation rate of implants in vivo was assessed over time.

#### 4.3. Histological analysis

Hematoxylin eosin (HE) stain, Masson stain of regenerated tissue were implemented to assess regeneration of tendon and muscle. The intermediate cross-section of the regenerated nerve was utilized for the Glycine silver (GS) staining procedure.

### 5. Statistics

All data were presented as Mean **±** Standard Deviation (SD) and analyzed by using Microsoft Excel software, t-test was employed to evaluate the difference between the groups; p < 0.05 indicates a statistically significant difference.

## Results

In our previous researches, the biomimetic materials have been applied as absorbable surgical sutures [He et al., 2025a], artificial tendons [He et al., 2025b], and hernia patches [He et al., 2025c]. Some pertinent figures from these preprints are adapted and presented concisely in the present research.

### 1. Implants fabrication and their properties

#### 1.1. Collagen thread (suture) and artificial implants

In this study, soluble collagen, the primary component, was utilized to fabricate collagen sutures (threads) that exhibit a range of color and diameter specifications. The collagen thread can function as an absorbable surgical sutures, it has been officially recognized by the CFDA (No. MZ17010224, No. WT17080974, No. WT19010144, No. WT19010145). In this study, different artificial implants were fabricated by weaving this collagen threads.

#### 1.2. Structure comparison between implants and native tissues

The collagen thread fabricated from soluble collagen is structurally composed of numerous tiny collagen fibers (Figure 2cd). This is corroborated by crushing a collagen thread monofilament after immersing it in physiological saline.

Artificial implants, such as hernia patches, artificial tendons, and nerve guide tubes (artificial nerve), are woven using collagen threads (Figure 2abi). Figure 2i illustrates a photograph of a hernia patch, its microscopic image reveals that it is constructed from collagen threads (Figure 2j). Figure 2ab depicts artificial tendons and artificial nerve, both of which are also constructed from collagen threads. Figure 2e shows a fresh rabbit Achilles tendon, Figure 2fg illustrates a bundle of natural tendons composed of numerous small collagen fibers. Masson staining of the rabbit tendon exhibits thick collagen fibers arranged in a highly densely organized manner (Figure 2h). Figure 2k depicts muscle tissue from the abdominal wall after immersion in formalin solution, Figure 2L showcases the densely arranged muscle fiber bundles that constitute the rabbit’s abdominal wall muscles. Figure 2m presents Masson staining of the muscle bundle, demonstrating the neatly arranged muscle fibers within the bundle. From this comparative analysis, it becomes evident that the collagen hernia patch or artificial tendon, artificial nerve effectively emulates the intricate microscopic structure of abdominal wall muscles or tendons, nerve epineurium respectively. Notably, the epineurium, the thickest connective tissue of the sciatic nerve, comprises 50-70% collagen. This protective layer (epineurium) shields the nerve from external compression by collagen fibers interwoven into a mesh-like structure.

**Figure 1.**
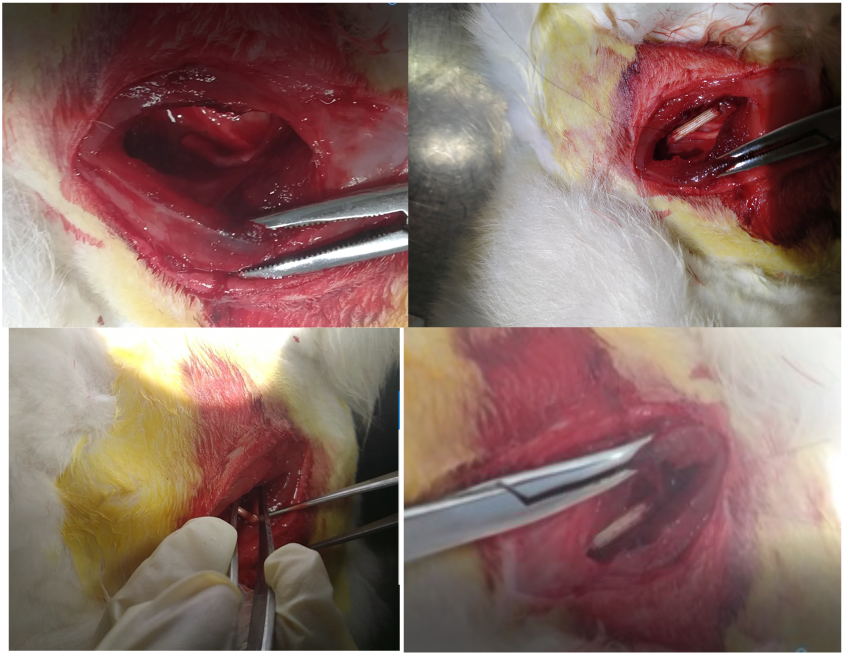
The surgical procedure of repairing 2-cm nerve defects utilizing a collagen nerve guide tube.

**Figure 2.**
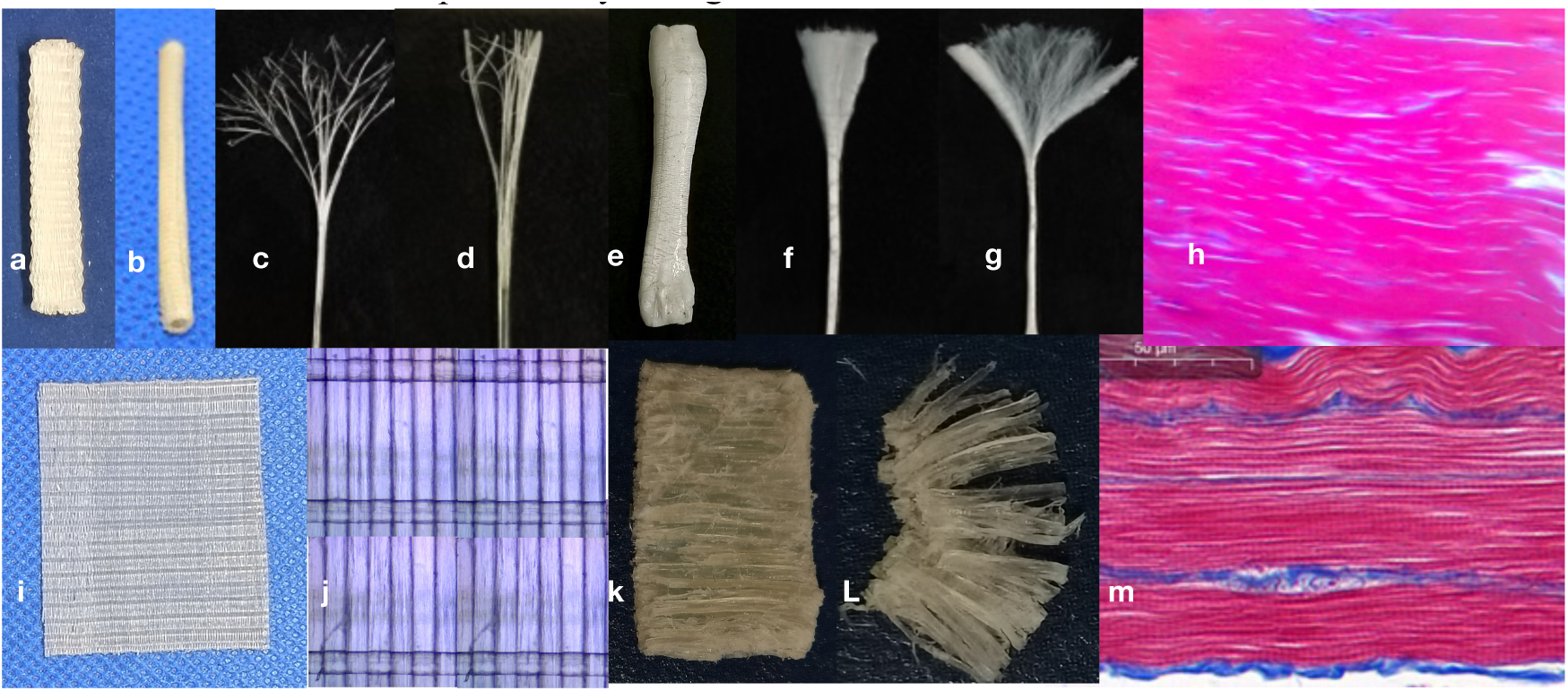
Comparison of artificial implants with autologous tissues. a: Artificial tendon; b: Artificial nerve; c and d: Structures of a collagen thread monofilament; e: Autologous tendon; f and g: Structures of a tendon bundle; h: Masson stain of tendon bundle; i: Hernia patch; j: Microscopic hernia patch; k and L: Structure of rabbit abdominal wall; m: Masson stain of rabbit abdominal wall (muscle).

Collagen, the fundamental structural protein of native tissue, is predominantly found in structural connective (loading) tissues. This collagen biomimetic material, characterized by exceptional mechanical strength, low antigenicity, and high affinity, can be utilized to emulate the majority of native tissues that contain collagen, including tendons and ligaments (70-85% collagen), the cornea (exceeding 90% collagen), blood vessels (primarily consisting of type I and III collagen, 50-70%), the gingiva (50-65%), visceral liver matrix (15-20%), the kidney matrix (approximately 10-15%), the gastrointestinal tract (around 25-30%), the fascia (60-80%), the uterus (40-50%), the trachea and bronchial walls (30-40%), the nerve epineurium (50-70%), the heart valve (50-60%), the gallbladder wall (35-40%), and the breast tissue (20-30%). By utilizing this biomimetic material technology, the repair or regeneration of these tissues can be effectively achieved.

#### 1.3. Tensile strength and suture strength of the implants

Mechanical strength of these implants are measured, and compared with native tissues as follows:

The tensile strength of artificial tendon, hernia patch, artificial nerve were shown in Figure 3. The tensile strength of dry artificial implants were significantly greater than native tendon, muscle, nerve respectively.

**Figure 3.**
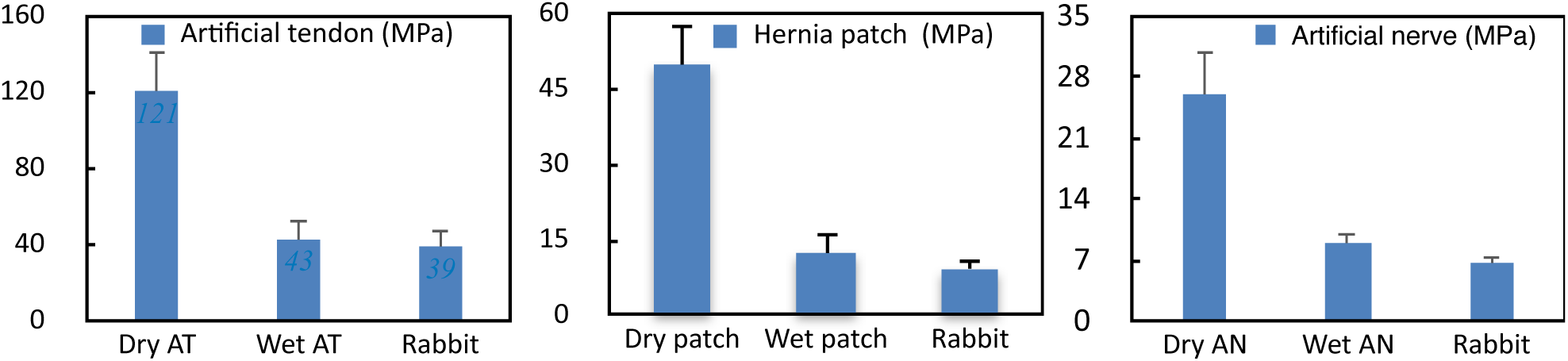
Comparison of mechanical strength between artificial implants and native tissues. AT: artificial tendon; AN: artificial nerve; Rabbit indicates native rabbit tissue respectively.

When these artificial implants were immersed in physiological saline for one hour, their strength decreased significantly. However, they still retain a comparable strength to native tissues, ensuring their ability to maintain strength in vivo (Figure 3). The mechanical strength of the wet artificial tendon and hernia patch was not significantly higher than that of native tendon and abdominal wall, respectively. Nevertheless, wet artificial nerve still exhibited significantly greater strength compared to native rabbit sciatic nerve (p< 0.05).

##### Suture Strength

Figure 4 illustrates the suture strength of artificial biomimetic implants and native tissue. Dry implants (tendon, patch, nerve) demonstrate significantly higher suture strength compared to fresh native tendon, abdominal wall, and sciatic nerve. Due to the hydrophilicity of collagen material, the mechanical strength of collagen implants experienced a rapid decline. Nevertheless, it retained its fundamental strength. Wet implants exhibit higher suture strength compared to native tissues without statistical difference (Figure 4).

**Figure 4.**
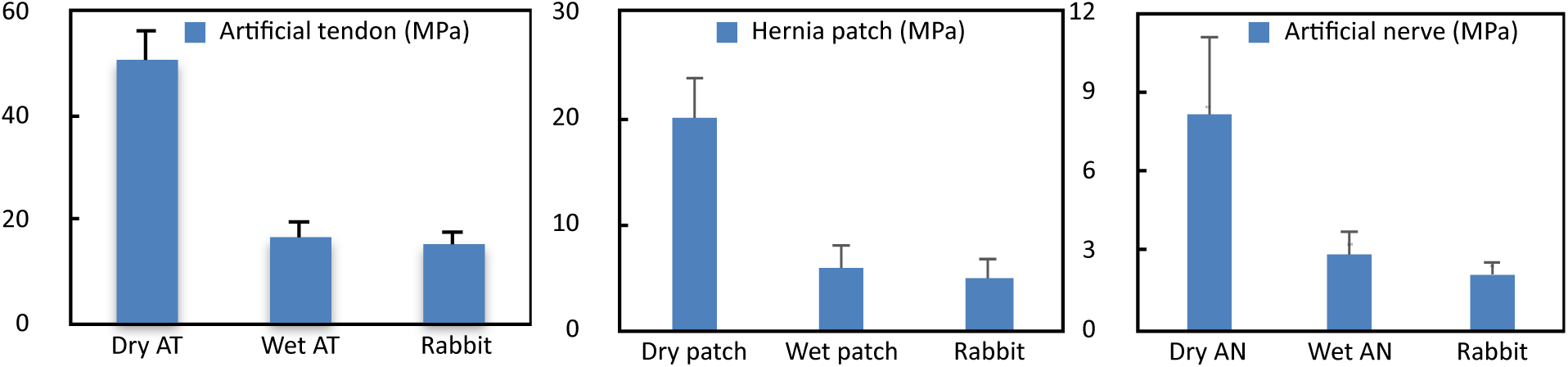
Comparison of suture strength between artificial implants and native tissues AT: artificial tendon; AN: artificial nerve; Rabbit indicates the rabbit native tissue respectively.

Based on the tensile and suture strengths of the implants, it is evident that their mechanical strength is sufficient to guarantee the in vivo safety and reliability after transplantation. Notably, throughout the entire animal experimental cycle, these artificial implants (tendon, hernia patch, nerve) exhibited no shedding, confirming that their broken ends can be securely sutured. It ensured the preservation of its mechanical strength in vivo until the regenerated tissue functions effectively.

The mechanical strength of these implants is directly related to their weaving technique, and their mechanical strength can be enhanced through intricate weaving.

As illustrated in Figures 3 and Figure 4, the mechanical strength of this biomimetic collagen material exhibit remarkable mechanical strength comparable to native functional tissue. Tendons, ligaments, and skeletal muscles are exceptionally robust tissues primarily dedicated to work, movement, and sports. Consequently, the exceptional mechanical strength of collagen biomimetic implants holds paramount significance, particularly during the initial phase of implantation. This underscores their potential to effectively address a diverse range of tissue functional regenerating requirements.

#### 1.4. Mechanical Strength of implants in vivo

These biomimetic collagen materials exhibit superior strength and integrity in vivo compared to absorbable synthetic materials of PLGA.

Following subcutaneous implantation, the tensile strengths of collagen sutures significantly exceeded those of the control sutures at 3 and 7 days (p < 0.05), as indicated in Figure 5 left. This is attributed to the fact that control sutures (PLGA) undergo degradation by hydrolysis, while collagen sutures are degraded by enzymes. Additionally, the immune system attacks and attempts to eliminate foreign matter-like PGA and PLA.

**Figure 5.**
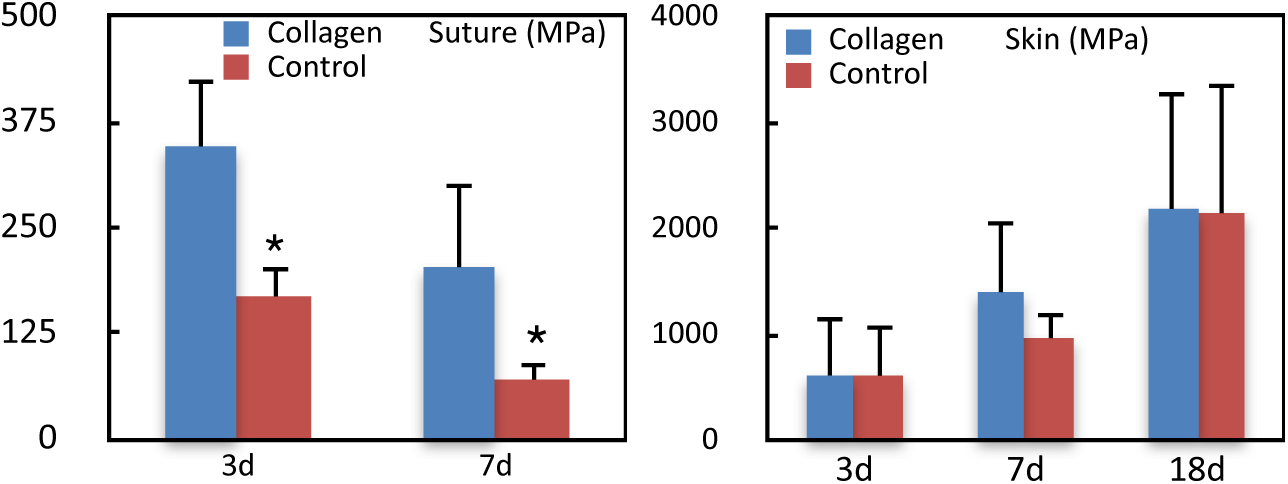
Left:Comparison of mechanical strength between test sutures and control sutures postoperatively. Right:Comparative mechanical strength of healed skin on test and control sides postoperatively.

Collagen sutures resulted in an increase in skin strength on the test side compared to the control side (PLGA suture) at 3d, 7d, and 18d points, as observed in Figure 5 right. Collagen sutures contribute more to wound closure than PLGA sutures. Control sutures (PLGA) may adversely affect rabbit skin wounds healing due to the formation of acidic by-products.

As depicted in Figure 6, these biomimetic collagen implants (artificial tendon/muscle/nerve) demonstrate exceptional mechanical strength before implantation, and maintain a substantial mechanical strength after implantation. Figure 6A illustrates the strength alterations in the regenerated tendon in vivo following transplantation. The artificial tendon initially exhibited a nonsignificant higher strength compared to the native tendon. However, this strength decreased over time, with the lowest value (25.1MPa) observed at the 8th week. Subsequently, the tendon underwent a rebound phase, reaching 31.3MPa in the 20th week, closely approximating the native tendon’s value of 39.1MPa, which represents 80% of the native tendon. Similarly, the collagen nerve tube and hernia patch exhibited a comparable pattern to the artificial tendon (Figure 6BC), with initial high strength followed by a reduction and subsequent rebound, ultimately reaching values that closely approximate the native tissue.

**Figure 6.**
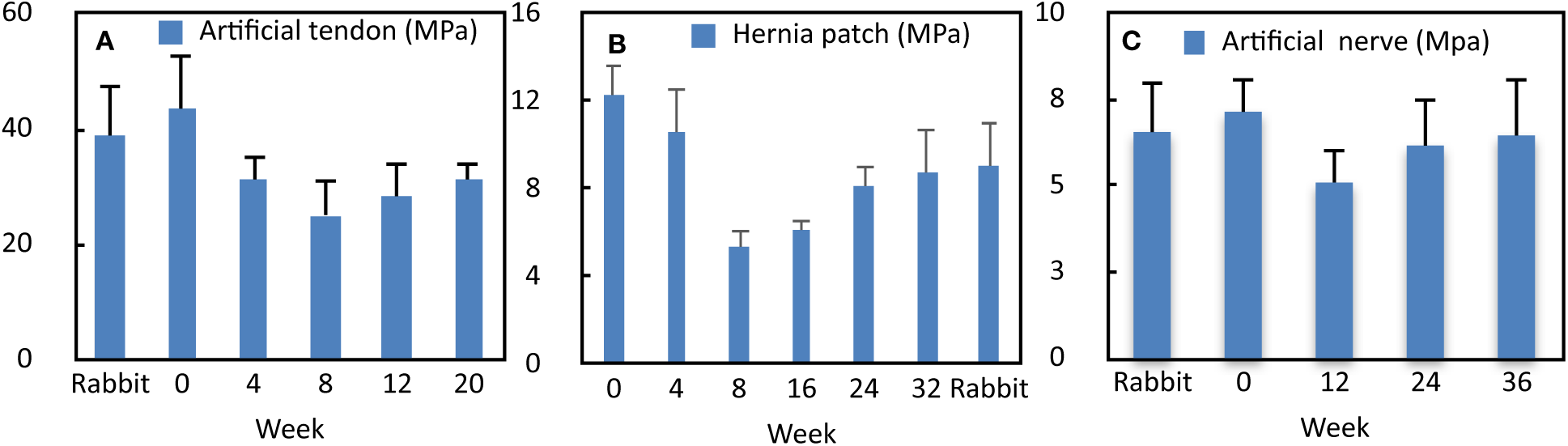
Mechanical strength of artificial implants after transplantation. Rabbit indicates the rabbit native tissue respectively.

All artificial collagen implants demonstrate exceptional mechanical strength, which is particularly pertinent for loading tissue, artificial tendons. These artificial tendons maintain their strength even at their lowest level 8 weeks after transplantation. This preservation of fundamental strength is crucial for the structural function and integrity of artificial implants. As observed in Figure 6, the strength of all collagen implants diminishes after transplantation. This suggests that various functional cells, including stem cells, invade and degrade the collagen implant materials. However, approximately 8 weeks post-surgery, the strength of the implants rebounds, indicating that the stem cells have transitioned into functional cells and secrete extracellular matrix (ECM), particularly collagen, to fortify the implants and establish regenerated functional tissues (tendon, muscle, nerve, etc.).

Figure 5 and Figure 6 illustrates the strength variation of collagen biomimetic implants after implantation, which closely correlates with cell activities in the local microenvironment. The strength of absorbable surgical sutures exhibits the most pronounced decrease in the skin, as the skin cells’ metabolism is the most rapid. Consequently, surgical sutures rapidly degrade and disappear. Similarly, the strength of the regenerating abdominal wall diminishes significantly due to the active abdominal muscular metabolism, then rebounds and recovers after approximately 8 weeks. Notably, the strength changes of nerves and tendons after implantation remain relatively stably slow, likely attributed to the slow metabolism of neurons and tendon tenocytes. Similar to absorbable sutures, all implants materials eventually degrade and disappear, contributing to the regeneration of functional tissues.

### 2. Animal experiments

#### 2.1. Overall appearance

Following surgical transplantation, the experimental animals were not administered immunosuppressive or anti-inflammatory drugs, except for intramuscular penicillin for consecutive three days. Throughout the experimental cycle, no discernible inflammation or immune responses were observed, no visible scar tissues, indicating that the collagen biomimetic materials exhibit low antigenicity and high affinity. Additionally, no implants experienced rupture or shedding, suggesting the stability, reliability, and safety of these implants. Throughout the experiments, regenerated abdominal wall (muscle) and nerve tissue were observed without adhesion. However, regenerated tendon tissue exhibited a notable adhesion to the surrounding tissue.

#### 2.2. Postoperative healing

As depicted in Figure 7 (top panel), collage sutures are employed to close skin wounds in rabbit models (the 1st row: black suture, 2nd rows: transparent suture), and in preliminary human application suturing an incision wound after the removal of a prosthetic from a woman’s breast (3rd row: transparent suture). Both kinds of collagen sutures exhibited satisfactory wound healing. Notably, the collagen sutures completely degraded by day 10 in rabbits, by day 9 in human. In human cases, the patient and the surgeon were satisfactory to the absorbable collagen sutures, smooth healing process companied with minimal inflammation and discomfort, and it is not necessary to remove sutures, which is particularly advantageous in surgery.

**Figure 7.**
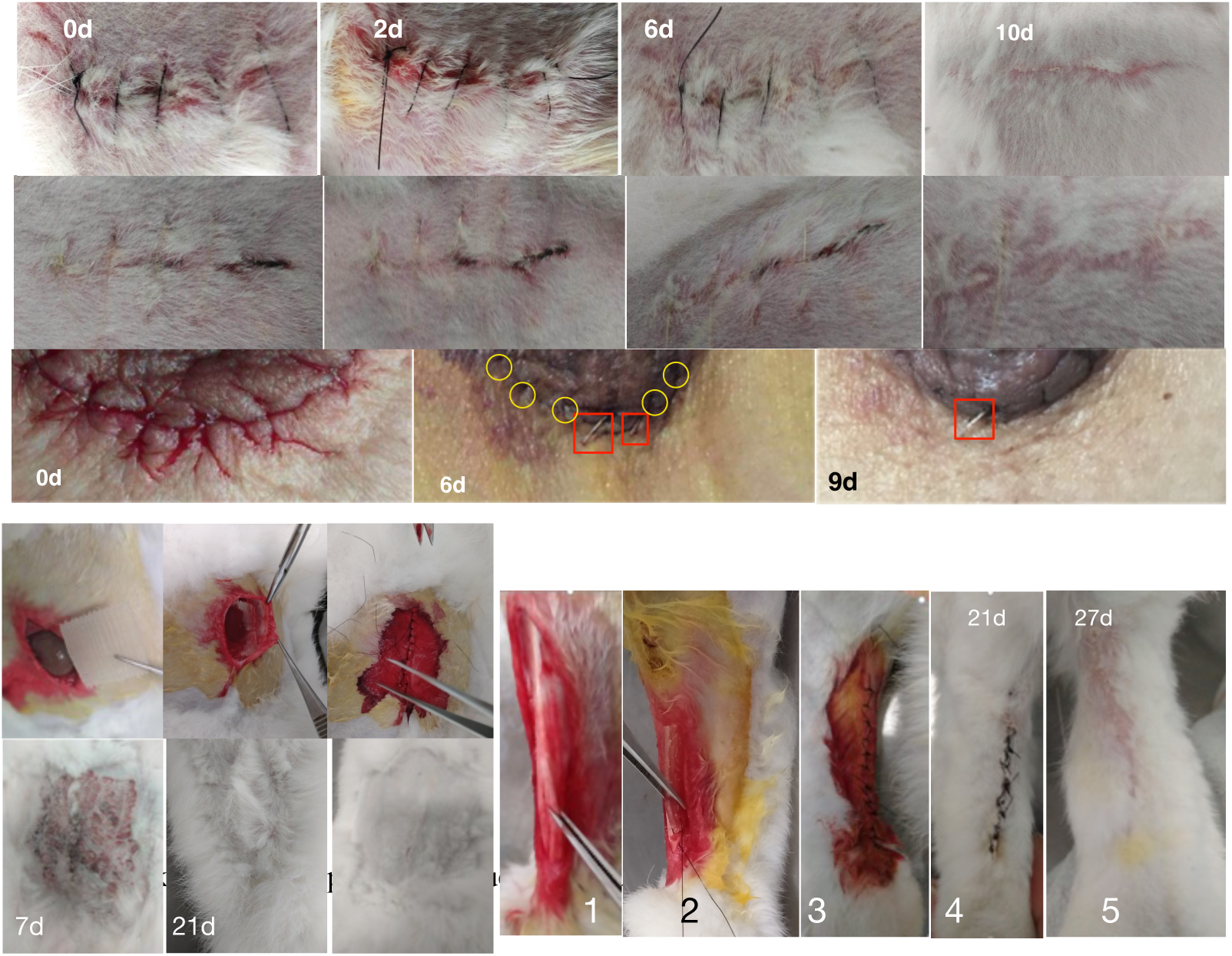
Top Panel: Collagen sutures were employed to close skin wounds. 1st and 2nd Row: Black and transparent sutures in a rabbit’s back. 3rd Row: Transparent sutures in a patient’s breast. Left Bottom Panel: Transplanting a collagen patch to repair a 3×3 cm VML. Right Bottom Panel: Transplanting an artificial tendon to repair a 2-cm tendon defect.

Figure 7 (right bottom) illustrates the transplantation of an artificial tendon to repair a 2 cm-long tendon defect. Figures 7-4 and 7-5 illustrate the skin wound healing very well. In the bottom left panel of Figure 7, following the implantation of a collagen hernia patch, the surgical incision on the skin healed satisfactorily within one week, and hair growth covered the surgical area after 3 weeks. The overall appearance was restored.

Tendons and abdominal muscles are situated just beneath the skin’s surface. Restoration of severe tendon or muscle damage significantly impacts skin wound healing. The satisfactory skin healing process is closely linked to the low immunogenicity and high affinity of the biomimetic implant material. If the implant material is non-biocompatible and highly immunogenic, the immune system perceives them as foreign substances, triggering the immune system’s defense mechanism. Macrophages attempt to engulf or encapsulate foreign materials. If degradation is not feasible, a fibrotic envelope forms. Additionally, cells release inflammatory factors (such as IL-1 and TNF-α), causing local tissue swelling, inflammation, fever, and pain. The immune system’s persistent efforts to eliminate the material result in chronic inflammation, potentially evolving into granulomas or fibrous tissue proliferation. If implanted materials damage the local tissue barrier, bacterial invasion occurs, leading to infections and suppuration in the skin or mucosal tissue. Some pathogens (such as Staphylococcus aureus) form a biological film on the surface of the implanted material, hindering the removal of infections by antibiotics. This infection exacerbates inflammation, further tissue damage, and potentially leads to wound ulceration. However, in this study, skin wounds heal remarkably well, even with only consecutively 3-day penicillin injection after the operation. This demonstrates the strong affinity of the material and its exceptionally low immunogenicity.

#### 2.3. Repairing volumetric muscle loss (VML)

Despite numerous efforts, the regeneration of volumetric muscle loss (VML) remains a significant challenge in the field of tissue engineering. However, Figure 8 depicts the successful regeneration of a 3×3 cm VML in the rabbit abdominal wall by using a collagen biomimetic hernia patch.

**Figure 8.**
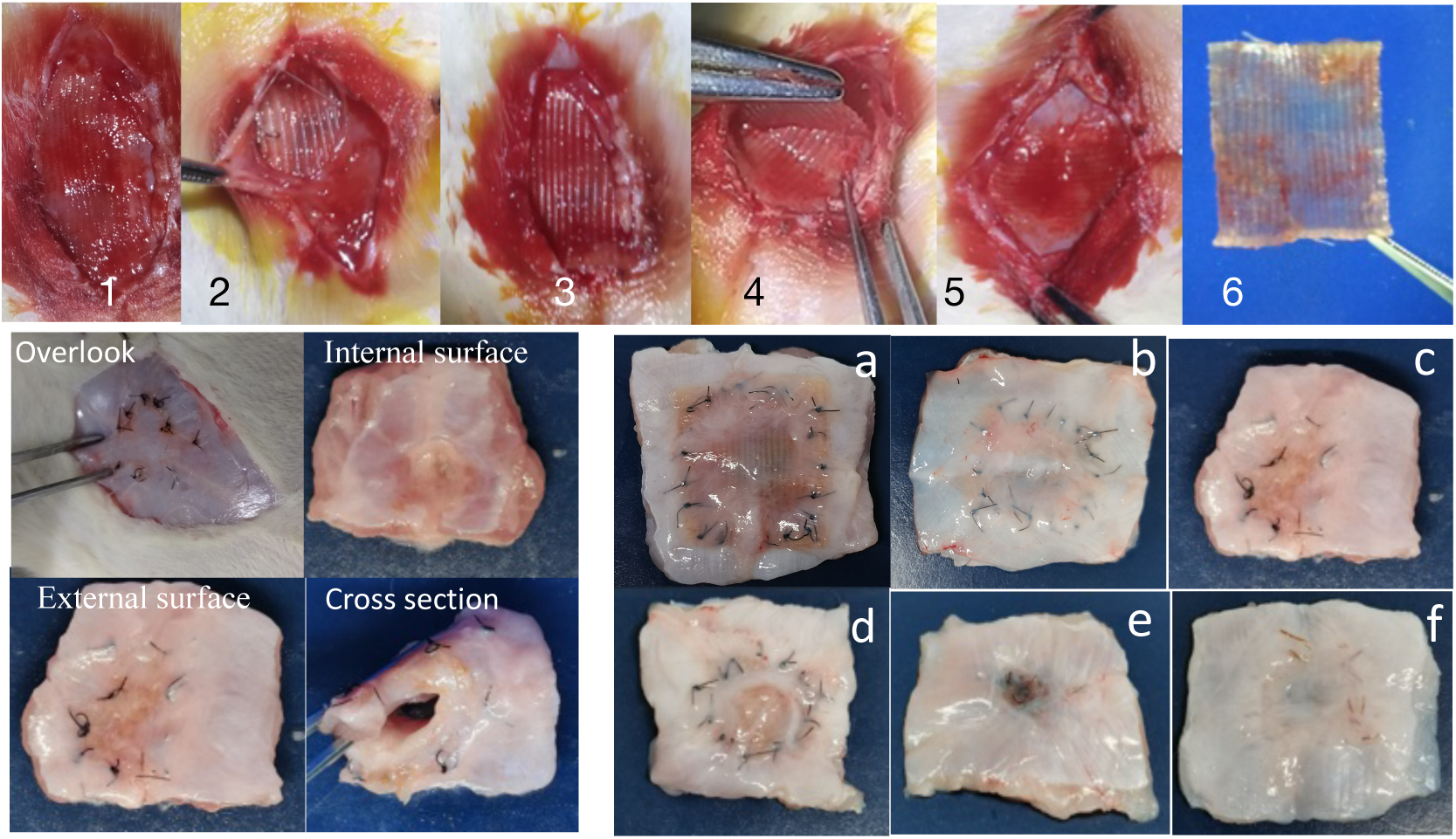
Transplantation of hernia patch and regeneration of abdominal wall Top panel: In vivo hernia patch 17days postoperative, left bottom panel: 16-week regenerated abdominal wall. Right bottom panel: regenerated abdominal walls in chronological order (abcde: nylon sutures of 4, 8, 16, 24, 32 weeks; f:collagen suture of 32 weeks).

In vivo evaluation of the implanted collagen hernia patch was conducted 17 days post-operation, as depicted in Figure 8 (top panel). The results revealed the formation of a three-tiered hierarchical structure. Newly formed tissues were observed growing on and beneath the collagen hernia patch, creating a sandwich-like structure. Specifically, a new membrane-like tissue covered the collagen patch (Figure 8-1, 8-2), and a new tissue layer also grew beneath the collagen patch (Figure 8-4), the covering tissue was removed without damaging the collagen patch (Figure 8-2, 8-3), the collagen patch can be separated from the underlying new tissue without damaging the patch or the new tissue (Figure 8-4,8-5,8-6), suggesting that collagen hernia patch can recruit numerous functional cells to form new tissues rather than recruiting fibroblast to form scar tissue. Structurally, the newly formed underlying tissue successfully isolates internal organs from the hernia patch, thereby inhibiting adhesion to internal organs. New vessels and blood were observed in underlying newly formed tissue (Figure 8-5), suggesting that collagen hernia patch can recruit capillaries or promote the formation of new vessels.

The 16-week regenerated abdominal wall muscle was excised and is depicted in Figure 8 (left bottom panel). Morphologically, it exhibits no significant distinction from the native abdominal wall.

The regenerated abdominal walls are presented in chronological order in Figure 8 (right bottom panel). As time passes, the nylon sutures gradually move from the surrounding wound anastomosis to the VML center, transforming the square wound defect into a circular one that shrinks over time. Around 32 weeks post-surgery, the sutures eventually concentrate at the center (Figure 8e). It suggests that the regenerated muscle tissues slowly grow inward, causing the nylon sutures to move inward as well. The hernia patch is mainly composed of collagen fibers, which can recruit numerous cells to degrade and utilize collagen materials as a source of ‘nutrients,’ thereby promoting cell growth and extension forward. This action effectively propels the nylon sutures inward.

There is no discernible boundary surrounding the anastomosis, nor any visible scar tissue, indicating that the muscle has been successfully regenerated and seamlessly integrated with autologous muscle tissue.

As previously illustrated in Figure 7, the biomimetic material does not induce inflammation of external skin wounds and facilitates the accelerated healing of external skin. Figure 8 further demonstrates that internal wounds heal successfully as well, attracting diverse functional cells that cover the collagen material. This observation reinforces the notion that the biomimetic material possesses properties of low immunogenicity and high affinity, underscoring its suitability as a potential medical regenerative material for tissue defect repair.

#### 2.4. Regenerated tendon

Until now, the regeneration of large tendon defects has been virtually intractable. However, this study presents a groundbreaking advancement in this field. Figure 9 demonstrates the tendon regenerating process after artificial tendon was implanted to repair the extensive defect.

**Figure 9.**
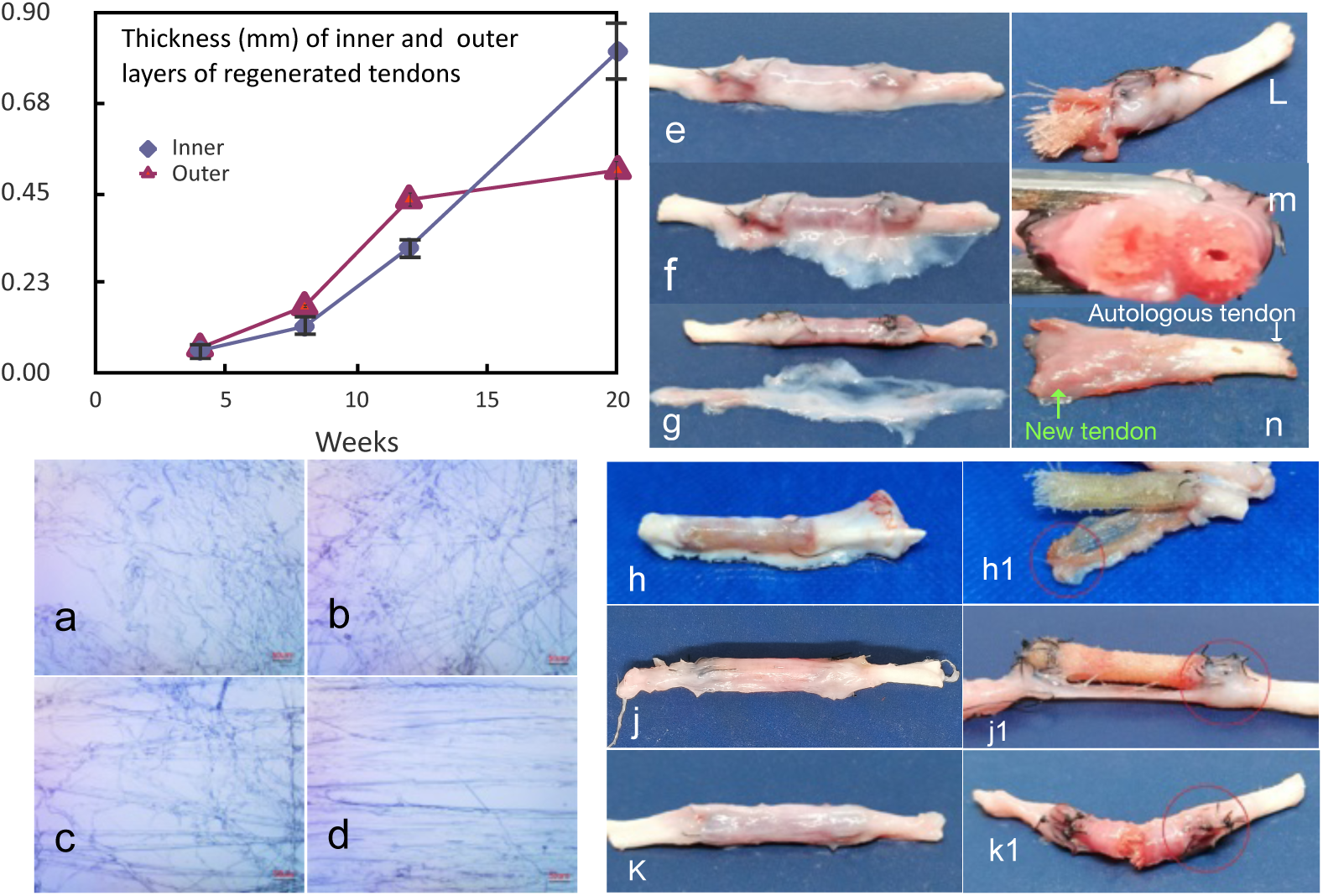
Tendon regeneration and regenerated tendons. Left A: Thickness changes of inner and outer layer of regenerated tendons; e,f,g: 20-week regenerated tendon. a,b,c,d: Morphological change of collagen fibers in outer layer; h,j,k: Regenerated tendons of 8,12, 20 weeks. h1,k1,j1:Disassembled newly formed tendons of 8,12, 20 weeks. Lmn: Half of the 20-week regenerated tendon. Green arrow: Newly formed tendon; White: Autologous tendon.

The 20-week regenerated tendon was excised and showed in Figure 9efg, a white outer layer of newly formed tissue covered the surface of the artificial tendon, the outer layer, is loose, can be easily separated by scalpel and scissors, then an inner dense tight layer revealed (Figure 9-g).

The outer layer of the white tissue, characterized by its loose texture, was predominantly composed of collagen fibers (Figure 9 abcd). These fibers underwent a gradual transformation from the outermost to the innermost regions, i.e., curving disorderly fibers were present at the outermost region (Figure 9a), followed by straight disorderly fibers (Figure 9bc), and finally, aligned orderly fibers parallel to the longitudinal axis of the autologous tendon were found at the innermost region (Figure 9d).

The thickness and state of the outer and inner layers also change with time goes (Figure 9A). Both layers progressively thicken over time. the outer layer is thicker than the inner layer before 12 week post surgery, however, after 12week post surgery, the inner layer is thicker than outer layer, and there is virtually no further thickening of the outer layer. The inner layer continues to thicken and develop a tougher texture, resembling that of native tendon.

Figure 9hjk depicts the regenerated tendons (without white loose outer layer) in chronological order, the color of the regenerated tendon changes gradually from blood-colored (red)(Figure 9h), transitions to a flesh-colored (pink)(Figure 9j), then shallowly white appearance (Figure 9k). The texture of the regenerated tendon were also gradually tougher and thicker, and resembled native tendon. Figure 9n depicted a regenerated flesh-colored tendon that seamlessly connected the natural autologous white tendon 20-week post surgery.

The strength of regenerated tendons increased over time. Upon removal of the nylon sutures, at 8 weeks, the newly formed tendon and the residual autologous tendon are readily separated (Figure 9 h1). However, at 12 weeks, the anastomosis is challenging to separate (Figure 9-j1). At 20 weeks, a thick, and tough inner layer has enveloped the artificial tendon, connection is firm and resilient, it is difficult to separate the connecting anastomosis (Figure 9-k1).

Similar to Figure 8, Figure 9 also illustrates the remarkable regenerative capabilities of this biomimetic material, characterized by low antigenicity, high affinity and high mechanical strength. Tendon regeneration presents greater challenges compared to muscle regeneration due to very limited functional cells, stem cells, and blood vessels in tendon region.

Muscle regeneration mechanism diverges significantly from that of tendon regeneration, which is facilitated by satellite cells locating within residual autologous muscles. Tendon regeneration, however, originates not from the residual autologous tendon but from the tissues surrounding the damaged tendon (paratendon tissue). This suggests that the biomimetic material possesses the ability to attract various functional cells to participate in regeneration.

This advanced material technology possesses substantial practical applications and can be employed to address a wide range of defect scenarios. For instance, the vascular wall is predominantly composed of collagen. Consequently, artificial vessels constructed using the biomimetic material would facilitate the regeneration of vessel tissue. This is particularly advantageous because small blood vessels (less than 4 mm) have not yet been successfully fabricated.

#### 2.5. Nerve regeneration

Numerous researchers have documented nerve regeneration, and nerve guide tubes (artificial nerve) have been commercialized by numerous companies. In this study, collagen biomimetic materials were also utilized to bridge a 2-cm defect in the sciatic nerve of rabbits. The results are as follows:

Figure 10 (top row) presents cross-sectional views of regenerated nerve tissue in chronological order. As time progresses, the tissue’s color becomes increasingly white, and the 36-week nerve is nearly identical to autologous nerve. Figure 10abc depicts 12-week regenerated tissue. After removing surrounding collagen materials, it was discovered that a newly regenerated nerve had bridged the 2 cm-defect (Figure 10b). Figure 10c provides a cross-sectional overall view of the 12-week regenerated nerve tissue, which is surrounded by numerous residual collagen fibers.

**Figure 10.**
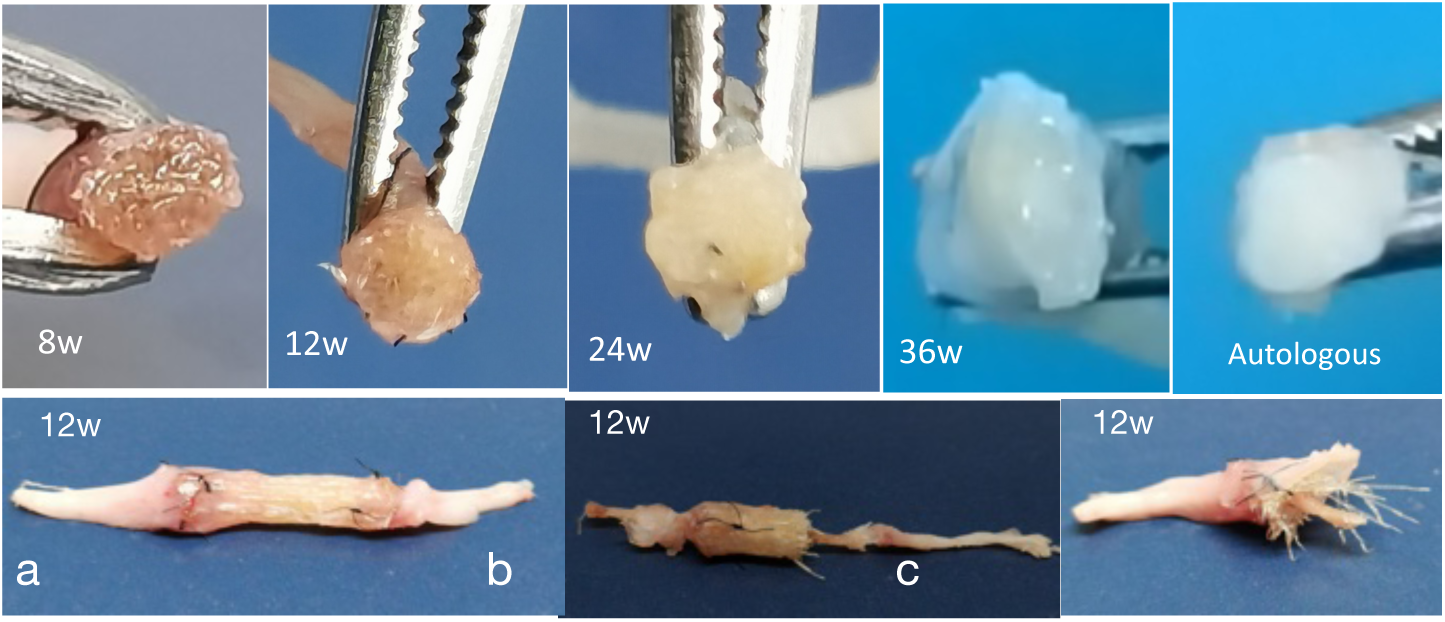
Regenerated nerve. Top row: Regenerated nerves in chronological order; Autologous: Rabbit sciatic nerve. Bottom row: 12-week regenerated nerve; a: Overlook; b: Regenerated nerve with half of collagen materials; c: Overlook of half regenerated nerve with residual collagen fibers.

Currently, commercially available medical products are designed to repair nerve defects. These materials, such as polylactic acid (PLA) and polyhexanoester (PCL), degrade gradually in the human body, effectively mitigating long-term foreign body reactions. However, this degraded acidic environment inevitably compromises nerve function. Early non-degradable materials like silicone and polytetrafluoroethylene (PTFE) have ever been utilized due to their commendable mechanical stability. Nevertheless, they may induce chronic inflammation or fibrosis reactions, necessitating surgical removal, therefore they are prohibited now.

The artificial nerve derived from collagen biomimetic material appears to be superior to other artificial nerve made from synthetic polymers like PLA and PCL. This is because the soluble collagen material facilitates nerve regeneration (nerve epinerium contains 50-70% collagen), resulting in accelerated and enhanced regeneration effects, conversely, the degradation byproducts of PLA, PLA are harmfully acidic.

The collagen biomimetic material exhibits versatility in tissue regeneration, encompassing both loading tissues like tendons and muscles, as well as non-loading tissues such as nerves. This broad application scope underscores the potential of biomimetic materials in tissue engineering.

### 3. Histological analysis

Histological analysis serves as a fundamental component in the investigation of pathological and structural characteristics of tissues. In this study, hematoxylin eosin (HE), Masson, and Gly Silver (GS) stains were employed to assess the regeneration of functional tissue. The staining outcomes of regenerated tendon, muscle, and nerve tissue are presented below:

#### 3.1. Tendon regeneration

Figure 11 presents a microscopic observation of regenerated tendon, obtained through direct observation and subsequent staining with HE and Masson stains.

**Figure 11.**
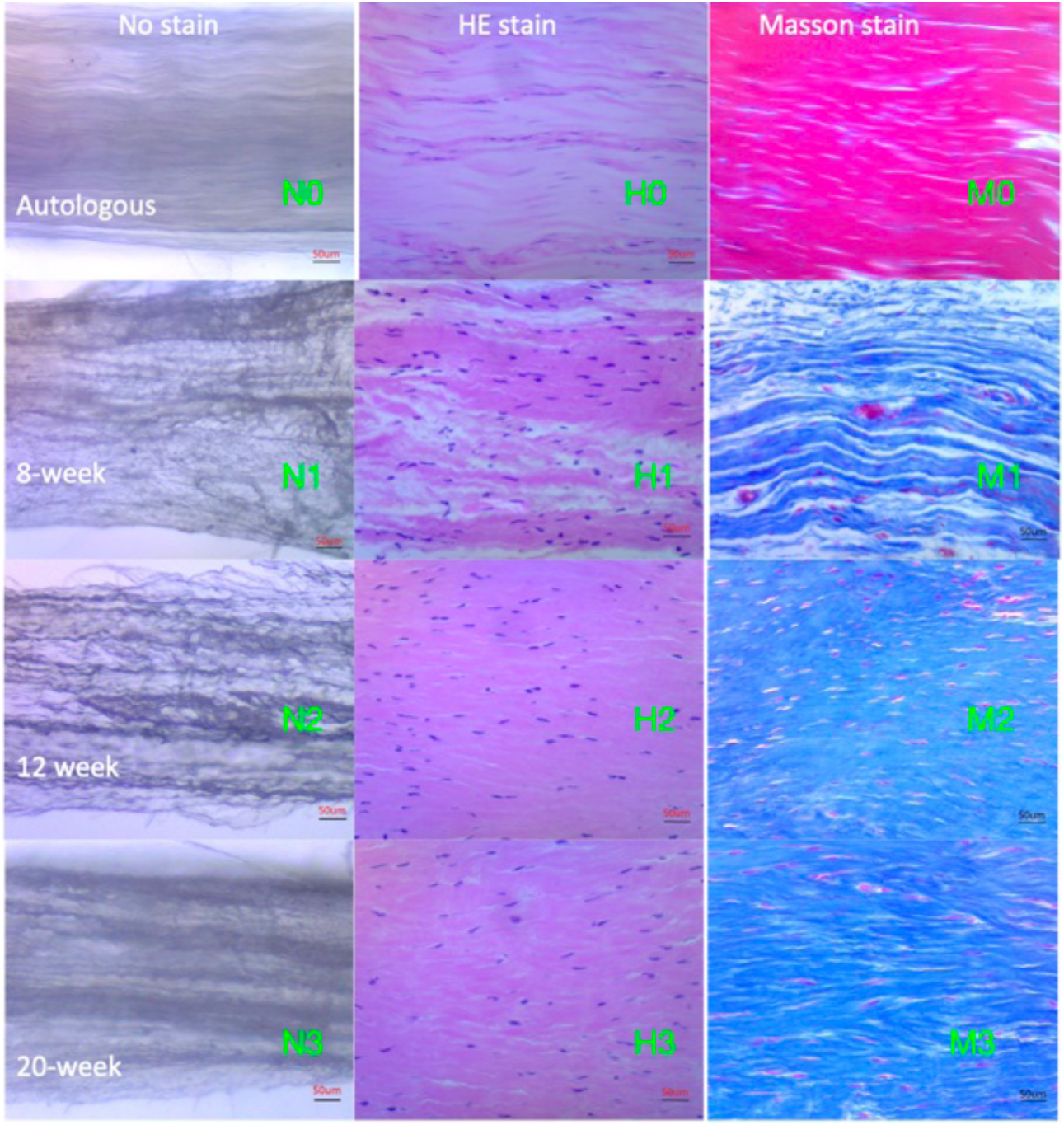
Microscopic structure of regenerated tendon and native tendon. Left column: regenerated tendon tissue without stain. Middle column: HE stain; right column: Masson stain; Top first row: native tendon, 2nd row: 8-week regenerated tendon. 3rd row: 12-week regenerated tendon; bottom row: 20-week tendon.

Figure 11N0 presents direct observations of native tendon, demonstrating densely arranged collagen fibers in a highly organized manner. These collagen fibers can be easily separated without causing fiber rupture. Figure 11N1,N2,N3 illustrates the newly formed collagen fiber tissues within regenerated tendons at 8 weeks, 12 weeks, and 20 weeks post-surgery respectively. As time progresses, the collagen fibers exhibit a gradual transition from sparse to dense. Initially, they are sparsely and slightly disordered, tightly bound (tangled). Disassembling these fibers without causing rupture proves challenging; however, this condition gradually improves over time, as the collagen fibers become denser, and collagen fiber disorder (tangling) degree decreases. This gradually allows for easy separation without rupture (Figure 11N2, N3). At 20 weeks, the regenerated tissue demonstrates dense collagen fibers, and the fibers become more orderly arranged, resembling autologous native tendons.

In contrast to direct observation, the HE stain can reveal the cell shape and quantity. Similarly, native tendon exhibits densely arranged collagen fibers (light staining) and a small number of flat fibroblasts (Figure 11 H0). From 8 to 20 weeks, the density of collagen fibers increases, similar to that observed directly, while the number of cells decreases over time. The cell shape transforms from oval to slender to flat (Figures 11 H1, H2, and H3). A comparable observation can be made in Masson stain (Figures 11 M0, M1, M2, and M3).

Figure 11 unequivocally demonstrates the mechanism and efficacy of tendon regeneration, simultaneously illustrating its gradually slow regeneration pace.

#### 3.2. Muscle regeneration

A hernia patch was utilized to repair a 3D muscle defect in the abdominal wall. It is observed from Figure 12 that the residual hernia patch was embedded within the regenerated tissue at 4-week and 8-week. Over time, the hernia patch underwent a gradual degradation and disappeared at 24 weeks. Muscle of Masson stain is red, as depicted in Figure 12 (Middle column), the red area of newly formed muscle tissue (Masson stain) exhibits a gradual increase over time, commencing approximately eight weeks after the surgical procedure.

**Figure 12.**
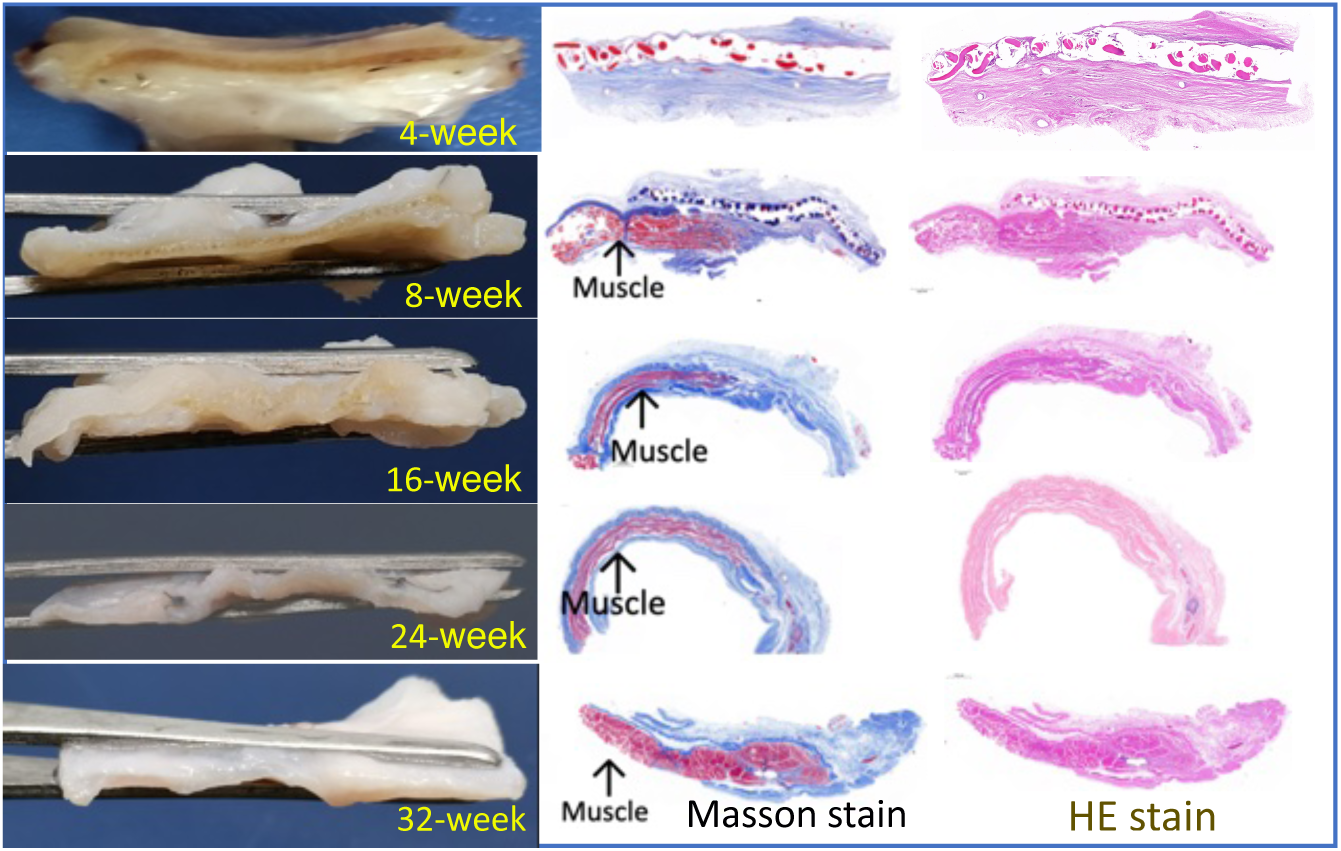
Cross sections of regenerated abdominal wall in chronological order (Left) and histological analysis (Right). Middle column: Masson stain (red muscle); Right column: HE stain (pink)

Figure 12 unequivocally demonstrates that muscle regeneration extends from the anastomosis site, thereby elucidating the robust regenerative capacity of satellite cells.

#### 3.3. Nerve regeneration

In this study, a range of staining techniques were employed to evaluate nerve tissue regeneration, with Gly Silver (GS) stain demonstrating exceptional efficacy in elucidating nerve structures. Figure 13A illustrates the GS stain of nerve tissue at 12-week, showcasing the small area of regenerated nerve surrounded by the majority of residual collagen fibers. This 12-week nerve is still in its primary stage, lacking a complete nerve tissue structure (Figure 13A). In contrast, the 24-week regenerated tissue exhibits an increased nerve area, accompanied by some residual collagen materials. The 24-week nerve demonstrates a clear nerve structure and some nervous tracts(Figure 13B). The 36-week regenerated tissue almost completely covers the defect, with minimal residual collagen materials. The 36-week nerve possesses an intricate structure with several nervous tracts (Figure 13C), similar to native sciatic nerve (Figure 13D). Notably, the native sciatic nerve possesses a highly intricate structure, comprising numerous neural tracts (Figure 13D).

**Figure 13.**
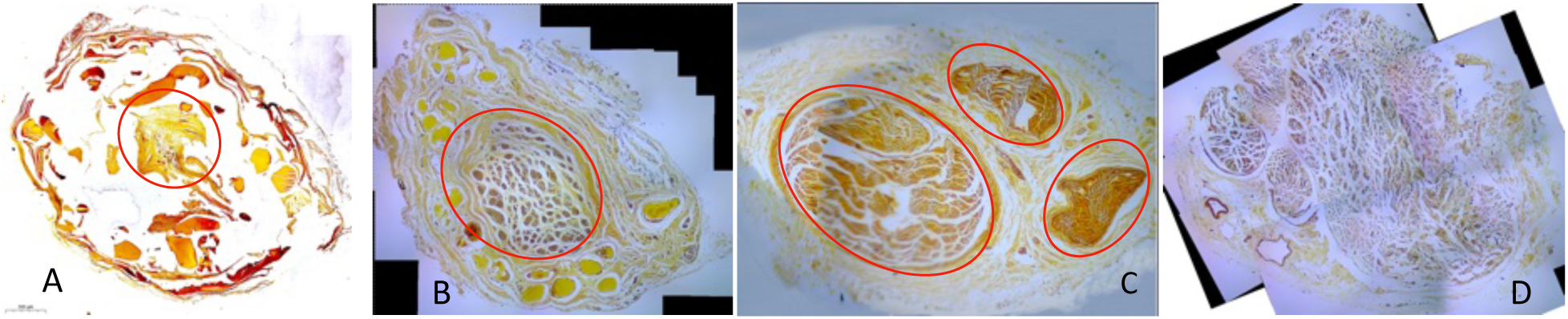
Gly Silver stain of 12-week (A), 24-week (B), 36-week(C) regenerated nerve and native sciatic nerve(D). The tissues surrounded by red circles are newly formed neural tissue.

### 4. In vivo degradation of implants

The biodegradation of scaffolds (implants) in vivo holds paramount significance for the advancement of regenerative medicine. Biodegradation is closely related to functional tissue regeneration. In this study, in vivo degradation rate of collagen absorbable surgical suture and collagen artificial implants (tendon, hernia patch, nerve) were calculated and showed in Figure 14, collagen surgical suture showed a statistically significantly (p<0.05) greater degradation rate compared to control suture (PLGA) at both 3 and 7 days after implantation. At 18 days, both of them degraded excessively, and there is no statistically significant difference between them (Fig. 14a).

**Figure 14.**
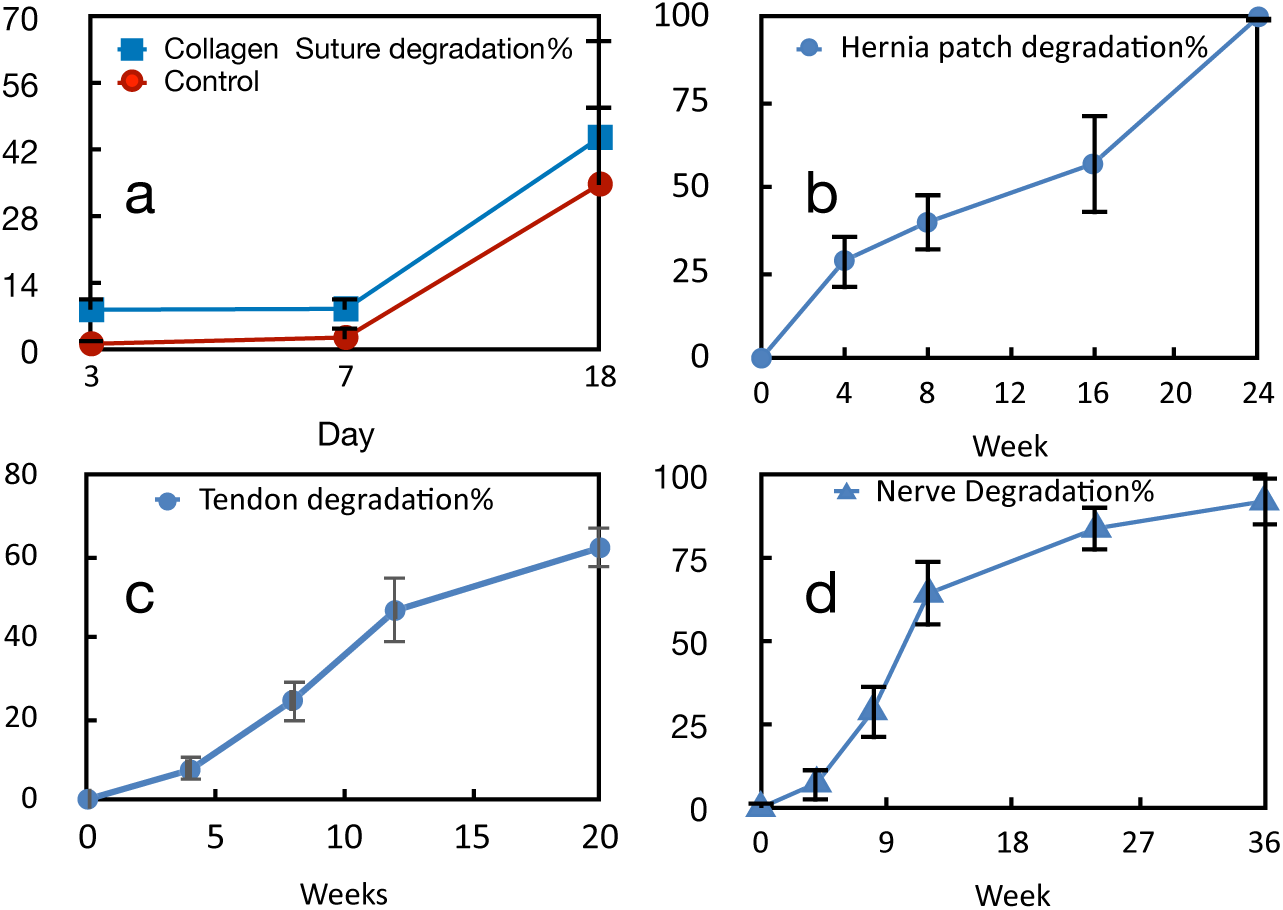
In Vivo Postoperative Implant Degradation.

The degradation rate of artificial tendon remains low during the initial four weeks, subsequently accelerating until the twelfth week, after which it decelerates (Fig. 14c). The degradation of artificial nerve exhibits a comparable pattern to that of tendon, as illustrated in Fig. 14d. However, collagen hernia patches demonstrate a distinct degradation pattern. During the first sixteen weeks, their degradation rate is relatively low, followed by a significant increase until complete degradation, rendering the patch invisible at twenty-four weeks (Fig. 14b).

The in vivo degradation processes of these biomimetic implants (Figure 14) correspond to the mechanical strength alterations of these implants in vivo (Figure 5, Figure 6).

As evident in Figure 14, collagen sutures in the skin undergo rapid degradation, with a 50% reduction observed 18 days post-surgery. The mechanical strength of the sutures decreased rapidly, simultaneously the strength of healing skin increased quickly, as shown in Figure 5.

In comparison to the degradation of sutures in the skin, degradation is a gradual process in regions such as tendons, muscles, and nerves. During the initial phase following implantation, cells began to infiltrate the artificial implants (hernia patches, tendons, and nerves), proliferate and simultaneously degrade the implants, resulting in a decrease in its mechanical strength. As the degradation of implants progresses, a significant number of cells accumulate, mature, and secrete a substantial amount of extracellular matrix (ECM). This ECM and cells form a functional structure, leading to an increase in the implant’s strength. The specific variation in the mechanical strength of implant is dependent on the type of target tissue.

Regarding the artificial tendon, the increasing collagen fiber density enhances its mechanical strength in the later phase, however, this fiber density hinders cell activity, leading to a decline in implant degradation. In contrast, during the later phase of muscle regeneration, muscle cells (muscle fibers) increase in number, and new vessels are formed. This further facilitates implant degradation. The substantially regularly arranged muscle fibers enhance the mechanical strength of the implant (hernia patch).

The varying pattern of artificial nerve is comparable to that of artificial tendon. Neurons (adult cells) are highly differentiated and exhibit minimal division or regeneration. Their primary function is to transmit and process electrical signals (action potentials) and chemical signals (neurotransmitters) rather than producing a substantial amount of anabolic substances. Energy primarily maintains electrochemical gradients (e.g., Na+/K+ pumps) and synaptic transmission. Consequently, their metabolic rate is low. However, if damage occurs in nerve tissue, axon regeneration of neurons commences. Axon regeneration and spread are highly energy-intensive processes that substantially accelerate implant degradation. As the axon moves forward, the intracellular energy (oxygen) reserve diminishes, necessitating increased resources for maintenance of the generated structure, thereby slowing down axon growth. Consequently, the degradation rate also decreases, and the mechanical strength of artificial nerves changes accordingly, similar to that of artificial tendons.

It is evident that the degradation of biomimetic implants exhibits a highly coordinated and synchronized relationship with the regeneration of targeted functional tissue. Functional regeneration is closely related to the microenvironment surrounding these implants. Collagen biomimetic materials can produce a niche microenvironment conducive to the growth and differentiation of stem cells (functional cells). This phenomenon underscores the exceptional characteristics of collagen biomimetic materials. If antigens and irritants within the implants induce immune or inflammatory reactions, the changing pattern of degradation and mechanical strength of implants would be completely different.

## Discussion

Tissue functional regeneration has been a longstanding pursuit of modern regenerative medicine.

This study has successfully developed a novel biomimetic material based on soluble collagen, addressing the longstanding challenge in the field of regenerative medicine. The novel regenerative material mimics the composition, structure, and function of native tissue, it can be fabricated to be biomimetic implants in place of autologous native tissue. Biomimetic implants augment the body’s inherent regenerative capabilities, unlocking the stem cell’s potential and enabling them to repair damaged defects, similar to autologous tissue.

The collagen biomimetic materials and their degradation products (a substantial number of biologically active fragments such as tripeptides and oligopeptides) establish a conducive environment for the growth and development of stem cells. The interactions between stem cells and the microenvironment ultimately facilitate cell differentiation and proliferation, resulting in the regeneration of functional tissue, such as muscles, tendons, and nerves in our research.

The three-tiered regulation mechanism, encompassing “material, degradation product, and microenvironment,” appears to be one of the fundamental reasons behind the successful regeneration of a substantial volume of functional tissues.

### Mechanisms of functional regeneration

Damage to the body that surpasses its inherent regenerative capacity is classified as a “critical defect” [Testa et al., 2021]. These collagen biomimetic implants facilitate functional regeneration of “critical defects” in diverse tissues. The internal regeneration mechanism can largely be attributed to the remarkable biomimetic properties of these implants in terms of composition, structure, and function.

Firstly, regarding composition, these collagen implants are derived from medical soluble collagen, which is devoid of cellular DNA, α-gal, and collagen telopeptides. Consequently, they can be recognized as “self” by the body’s immune system, thereby significantly reducing inflammatory and immune reactions. This minimizes the likelihood of scarring or fibrosis [Badylak et al., 2009]. The collagen material can induce macrophages to be M2 (repair-promoting) phenotype, releasing anti-inflammatory factors such as TGF-β and IL-10, thereby reducing inflammation. Collagen effectively synergistically combines with various active molecules, such as integrins (α2β1, α5β1) and growth factors (IGF-1, HGF, VEGF), to establish a localized microenvironment with high concentrations of bioactive molecules. This microenvironment is conducive to stem cell proliferation and differentiation. Collagen scaffolds and its degradation byproducts also play an important role in recruiting stem cells to the damaged site, and guiding stem cells to migrate, proliferate, and differentiate into target cells through CXCR4/SDF-1 pathway, FAK/ MAPK signaling pathway, or Notch, Wnt, Smad signaling pathways [Engler et al., 2006; Guilak et al., 2009; Li et al., 2004; Levenberg et al., 2005; Schwartz & Ginsberg, 2002; Zammit et al., 2004].

Secondly, the collagen fiber arrangement of the biomimetic implants closely resembles the natural fiber structure of body tissue. such as tendon, ligament or muscle tissue. This topological structure directs the activities of stem cells in a specific manner, facilitating their expansion, migration, proliferation, and differentiation. Consequently, a well-structured tissue is reconstructed [Choi et al., 2008; Angelina et al.,2018; Hochleitner et al., 2018; Yin et al., 2010; Zheng et al., 2017]. On the other hand, our cross-linking approach greatly improves the mechanical strength of the biomimetic implants. These biomimetic implant materials can not only substantially support the implant structure at the initial phase but also can keep its strength for tissue remodeling until implant degradation and complete tissue regeneration. Furthermore, effective cross-linking also delays the activity of collagenase and matrix metalloproteinase (MMP) in degrading collagen implants. Consequently, the degradation of the collagen implants can be synchronized with tissue reconstruction and the functional regeneration of new tissue [O’Brien, 2011], which ensure the safety and reliability of implant transplantation throughout the entire regeneration process.

Lastly, these collagen biomimetic implants not only maintain the mechanical strength, the transplantation of these biomimetic implants also induces in vivo revascularization. Collagen and its degradation products stimulate the formation of new blood vessels, facilitating tissue repair and functional regeneration [Davis and Camarillo, 1995; Ruhrberg et al., 2002]. As mentioned before, M2 macrophage phenotype induced by collagen materials secretes VEGF, FGF, attracting endothelial cells and form new capillaries (diameter 10–50 µm), which ensure the blood supply of regenerated tissue [Murray, 2017: Levenberg et al., 2005]. Short peptide (<10 kDa) from collagen degradation improves the stability of new blood vessels through the VEGF/ANG-1 signaling pathway and prevents ischemic collapse, new vessels play a critical role in regeneration of large tissue [Wang et al., 2009]. Besides vessel, neural existence is also a pivotal factor to tissue reconstruct especially muscle regeneration. The collagen fiber structure closely resembles the mechanical microenvironment of structural tissues, offering physical cues that promote the growth of nerve axons and facilitate the reconstruction of nerve-muscle junctions [Jessen and Mirsky, 2016; Kjaer, 2004; Chen et al., 2007]. As previously discussed, VEGF released by M2 macrophages promotes angiogenesis, VEGF also plays a crucial role in releasing neurotrophic factors, which facilitate the integration of neurons with newly regenerated tissue [Gonzalez et al., 2011].

These biomimetic implants derived from soluble collagen exhibit remarkable properties, including exceptional strength, high affinity, low antigenicity, and high nutritional value. Collagen and its degradation products can create a biological microenvironment that facilitates the growth of stem cells and functional cells. Consequently, the biomimetic scaffold participates in the body’s metabolism and gradually integrates into the body’s structure. This transformation elevates the biomimetic scaffold from a non-living implant to a living regenerated tissue.

### Implant integrity, strength and function in vivo

The biomimetic materials demonstrates its extensive regenerative capabilities in tissue engineering. It is crucial to elucidate the mechanisms by which biomimetic collagen materials preserve their integrity, strength, and functionality in vivo. This analysis seeks to offer an alternative perspective on the regeneration mechanism.

Firstly, it is because of collagen itself not synthetic degradable polymer, as mentioned previously, synthetic degradable polymer scaffold would collapse quickly in vivo due to hydrolysis; secondly because of effective cross linking, the collagen biomimetic material delays enzymatic degradation (e.g., collagenase, MMPs) within a 6-12 month timeframe [Parenteau-Bareil et al., 2010], it is enough for collagen scaffolds to recruit numerous stem cells, which entails the biomimetic material similar to autologous tissue, its degradation rate aligns with the reshaping rate of the newly formed tissue, preventing biomimetic materials collapse at too fast a rate and fibrosis at too slow a rate. In this way, host functional cells (including stem cells) gradually replace the implant material and maintaining the scaffold’s integrity [San Antonio et al., 2020; Ferrara & Davis-Smyth, 1997]. Simultaneously, this cross-linking also enhances the mechanical strength of the biomimetic material, it can withstand physiological loads, such as tendon stretching forces, maintain its elastic modulus, and prevent postoperative tearing [Holzapfel, 2001]. During the degradation process of collagen materials, new intermolecular cross-linking (in vivo cross linking) or recombination may occur, temporarily strengthening the remaining structure [Hematti, 2018]. Additionally, collagen’s enhanced integration with host tissue promotes the growth of tissue cells on the implant and the secretion of ECM (e.g., collagen)[Brown and Philips, 2007], this gradual accumulation helps to keep the implant’s integrity and enhances its mechanical properties [Butler et al., 2008].

The complete functionality of regenerated tissue is derived from functional cells and their secreting active molecules or extracellular matrix (ECM). For instance, tendon functions due to collagen fibers secreted by tenocytes, while muscle function is enabled by muscle fibers derived from satellite cells. Exogenous cells are not implanted in our researches, the partial functionality primarily stems from the structural integrity and mechanical strength of the biomimetic implant during the initial phase following artificial tendon (muscle etc.) transplantation, the ordered collagen fiber arrangement of this material mimics the parallel fiber structure of natural tendons (or muscles etc.), maintaining directional mechanical properties. Subsequently as mentioned before, the biomimetic material can combine various active molecules, recruiting cells, promoting cell proliferation, vascularization, innervation, and ECM deposition [Roberts et al., 1986]; additionally, the biomimetic structures guide stem cell migration, differentiate and matrix secretion along the mechanical direction to form tissue-like structures, such as tendon-like structures restoring tensile strength and partially restoring the sliding function of muscles [Dunn & Silver, 1983; Zhao et al., 2010, Zhang & Wang, 2010], thus gradually restoring full tissue functions.

Generally, the body’s immune system, blood vessels, nervous system, stem cells, and other components collaborate to preserve the integrity, mechanical strength, and function of the biomimetic implant in vivo. This collaboration and coordination of immune regulation, stem cell recruitment, signal transmission, vascularization, and innervation facilitate tissue remodeling and functional tissue regeneration. Notably, the immune system not only refrains from attacking the biomimetic materials framework, but also it actively participates in functional regeneration, thereby preserving its initial integrity. For instance, the M2 phenotype, induced by collagen material, releases anti-inflammatory factors such as TGF-β and IL-10, thereby reducing inflammatory damage. Additionally, the M2 phenotype releases VEGF to promote the formation of new vessels.

### Comparison with other implant materials

As previously discussed, conventional implant materials, such as synthetic polymers, metals, ceramics, and dECM, are not optimal for tissue engineering. The body’s defense mechanism impedes functional regeneration. Even in instances of mild inflammation or a moderate immune response, regenerated tissue frequently fibroses, resulting in scar tissue formation [Chen et al., 2009].

Consequently, numerous research endeavors have focused on utilizing natural proteins derived from the human system. However, it is crucial to acknowledge that macromolecules within our bodies, such as fibronectin, laminin, and hyaluronic acid, are also not suitable as the primary component of scaffolds in place of collagen. Adhesive protein (Fibronectin), a glycoprotein, plays a pivotal role in cell adhesion and migration. However, it lacks the capability to form highly structural organized fibers for load-bearing [Gautieri et al, 2011]. Hyaluronic Acid (HA) is a macromolecular polysaccharide primarily responsible for lubrication and hydration. It can absorb a substantial amount of water and maintain tissue volume, it lacks tensile strength, can not support a structure. Laminin, a structural protein, is predominantly found in the basal membrane. Its primary functions encompass cell adhesion and signal conduction, rather than serving as the primary load-bearing structure [Fritz, 2008; Shoulders and Rainers, 2009]. Collagen fibers exhibit exceptional tensile strength of 50-80 MPa, possessing a high elastic modulus, which enables them to withstand significant tensile forces without deformation..

Collagen molecules establish covalent cross-links through lysine residues, including pyridinoline and deoxypyridinoline, which significantly enhances its anti-enzymatic properties and mechanical stability. In contrast, fibrin, hyaluronic acid, and other biomolecules lack a comparable covalent cross-linking mechanism, resulting in structural stability that is considerably inferior to that of collagen [Eyre et al., 1988].

Collagen has its unique molecular structure, biomechanical properties, tissue-specific distribution, chemical stability, and evolutionary conservation. These characteristics collectively determine its irreplaceable role in preserving the structural integrity of mammals. While adhesion proteins, hyaluronic acid, and other components contribute to the extracellular matrix, they cannot assume the structural and mechanical support functions that collagen performs.

In comparison to other materials derived from the human body, soluble collagen emerges as the highly significant and viable option for fabricating exceptional scaffolds in tissue engineering. This is also the reason why biomimetic materials based on soluble collagen can functionally regenerate tissues successfully.

### Collagen biomimetic material and stem cells

As previously discussed, collagen is the most effective macromolecule capable of serving as the primary component of scaffolds in tissue engineering. Collagen’s multifaceted role extends beyond its structural function, encompassing a diverse range of aspects. It collaborates with various active molecules, growth factors, facilitates intercellular communication, and transmits signals from the cellular level to the tissue level. Consequently, collagen holds a crucial “nutritional” value for organisms.

In this study, soluble collagen-based implants exhibited successful regeneration of tendons, muscles, and nerves by supporting stem cells or functional cells. This is primarily attributed to the five pivotal roles of collagen material.

The first one, collagen creates biomimetic niche microenvironment in vivo to control stem cell behavior by providing critical structural and biochemical cues. As mentioned before, collagen’s triple-helix structure, tunable mechanical properties of 3D collagen scaffold mimic tissue-specific microenvironment, facilitate spatial cell distribution and tissue-like organization. Collagen also supports nutrient and oxygen diffusion, enhancing MSC proliferation [Chan & Leong, 2008], collagen and its degradation byproducts combine various growth factors, active molecules, promoting survival, proliferation and differentiation of stem cells [Tibbitt & Anseth, 2009]. Collagen type, concentration, and cross-linking degree can substantially influence the fate of stem cells, encompassing self-renewal and differentiation [Gershlak et al., 2017].

Secondly, collagen promote stem cell adhesion & proliferation, which are prerequisites for effective tissue regeneration. RGD-mediated integrin binding significantly improves stem cell adhesion, reducing apoptosis. Collagen gels have been shown to support adhesion of adipose-derived stem cells (ADSCs) and embryonic stem cells (ESCs) [Tibbitt & Anseth, 2009]. Collagen scaffolds ensure efficient nutrient and oxygen exchange, enhance MSC expansion and proliferation [Rustad et al., 2010].

Thirdly, collagen scaffolds guide stem cell differentiation through tailored mechanical and biochemical cues, addressing diverse tissue regeneration needs. Stiff collagen scaffolds induce MSC differentiation into osteoblasts, while soft collagen gels favor chondrogenic or neurogenic differentiation [Bian et al., 2013]. Collagen scaffolds can combine with growth factors (e.g., TGF-β1, BMP-2, VEGF), thereby regulating their survival, migration, and differentiation [Kim et al., 2011]. BMP-2-loaded collagen scaffolds promote MSC osteogenesis for bone defect repair [Dawson et al., 2008]. Aligned collagen fibers direct cell morphology and differentiation, oriented collagen scaffolds guide neural stem cells (NSCs) to form organized neural tissues (Lu et al., 2011).

Fourthly, collagen can function as cell delivery vehicle, in vivo stem cell delivery faces challenges such as low survival rates and poor localization. Collagen provides a physical barrier, shielding stem cells from mechanical stress and immune responses [Lee et al., 2011]. In our muscle regeneration experiment, collagen biomimetic implants (hernia patches) effectively deliver satellite cells to bridge the significant defect. Collagen gels also immobilize stem cells at target injury sites, preventing dispersion. MSCs delivered via collagen scaffolds enhance bone defect repair [Tabata, 2009]. By modulating scaffold degradation rates (e.g., via EDC or glutaraldehyde cross linking), collagen enables sustained stem cell release, prolonging therapeutic effects. Cross-linked collagen scaffolds achieve gradual MSC release, improving cardiac function in myocardial infarction models [Kim et al., 2013].

Fifthly, collagen scaffolds also support functional cells (e.g., osteoblasts, endothelial cells) to promote tissue functionality, collagen mimics native ECM, promoting matrix secretion by functional cells. For instance, fibroblasts in collagen scaffolds produce higher levels of collagen, aiding skin regeneration [Geckil et al., 2010]. Collagen facilitates integration of functional cells with host tissues. Endothelial cells in collagen matrices form vascular-like structures, integrating with host vasculature [Glowacki & Mizuno, 2008]. Collagen facilitates the physiological activities of functional cells, thereby accelerating the restoration of functional tissue.

The biomimetic implant materials based on soluble collagen exerts a pivotal role in supporting the cellular functions of stem cells and functional cells, thereby facilitating the functional regeneration of a wide range of tissues. In a word, its exceptional biocompatibility, low antigenicity, biodegradability, mechanical strength, and bioactivity contribute to this outcome.

### Structural Biomimetics in Tissue Engineering

The scaffold structure plays a pivotal role in determining the success of tissue engineering endeavors. Optimal scaffolds are biomechanical and biocompatible, facilitating cell attachment, proliferation, differentiation, and guide cellular behavior. However, these requirements often present conflicting. For instance, dECM scaffolds offer tissue-specific intricate architecture but compromise bioactivity and immunogenicity. Conversely, simple scaffolds of collagen sponge (or gel) exhibit high affinity, enhancing nutrient diffusion but compromising mechanical strength, our biomimetic scaffold addresses the paradox and achieves a balance between structural integrity and functional efficacy,

It appears that a scaffold with a meticulously structured design is preferable to a scaffold with a simple design, provided that the same materials are employed. However, a finding from our study indicates that this assumption may not always hold true. The structural biomimetics employed in our study for muscle regeneration exhibit intriguing results. Normal 2 mm abdominal walls were successfully regenerated by using a single-layer 0.4 mm collagen hernia patches. There exists a notable disparity between 0.4 mm and 2 mm; a collagen patch may not accurately simulate the abdominal wall structure. To address this limitation, we employed a 4-layer collagen patch (approximately 1.6 mm thick), which is more closely aligned with the anatomical structure of the native abdominal wall. Surprisingly, the outcome was not as favorable as that achieved with a single-layer 0.4 mm collagen patch. It appears that the structure is not beneficial to skeletal muscle cells (muscle fibers); the detailed mechanism should be further investigated.

The scaffold structure exhibits a correlation with other characteristics of the scaffold. Together, these characteristics influence regeneration efficiency and the ultimate outcome, necessitating a detailed analysis.

There are primarily three types of scaffold structures: complex, simple, and intermediate. Firstly, dECM scaffolds, mimicking intricate native tissues, have ever been regarded the best scaffolds, however, they are subtly different from native tissue in chemical composition, surface topography, degradation kinetics, and mechanical properties (e.g., stiffness) [Turner & Badylak 2012]. Their regeneration efficiency has been significantly hindered primarily by trace residual antigens (e.g., cellular DNA, Galα1-3Gal epitopes, telopeptide) [Wong & Griffiths 2014; O’Brien 2011], dECM scaffolds also lack the bioactive cues (e.g., FGF, HGF, VEGF, IGF-1 and other bioactive molecules) necessary for stem cell activation and vascularization [Sicari et al., 2014], which were removed by decellularization [Lutolf & Hubbell, 2005], impairing angiogenesis and cell differentiation [Keane & Badylak, 2015]. Dense or overly complex structures of dECM may restrict cell infiltration and nutrient diffusion [Hollister, 2005]. Stiffness and topography of dECM scaffold inhibit stem cell differentiation [Engler et al., 2006]. In contrast, simple scaffolds of natural polymers (e.g. hydrogels and sponges from collagen and gelatin) exhibit high biocompatibility and low immunogenicity, their soft, loose, and porous structures facilitate cell invasion, enabling them to secrete and remodel their own ECM. Collagen also can capture growth factors, adapt to the local microenvironment, and demonstrate remarkable efficacy in specific applications [Badylak, 2014; Wolf, et al. 2012]. However, their limited mechanical strength restricts their further utilization, particularly in load-bearing tissues. Thirdly, structure of synthetic scaffolds (e.g., PCL, PLGA) is highly tunable (e.g., pore size, mechanical properties) but lack biocompatibility and native biochemical cues (e.g., growth factors, adhesion molecules)[Lutolf & Hubbell 2005], and its acidic degradation products may cause inflammation, restricting their wide application.

Consequently, it becomes evident that the optimal structure should possess the capacity to accommodate a sufficient number of stem cells or functional cells.

Our biomimetic scaffold possesses advantages of above 3 types of scaffolds, striking a delicate balance between structural complexity, bioactivity, mechanical properties, and immunogenicity, facilitating cellular remodeling and integration. The scaffold degradation is controlled by in vivo niche environment (pH, temperature, or enzyme levels etc.), dynamically adapt to tissue remodeling and match tissue regeneration [Rice et al., 2013].

Future scaffold should be multi-scale design, integrate macro-(mechanical support), micro-(porosity, fiber alignment), and nanoscale (surface chemistry) features [Moroni et al., 2018], and computational modeling can predict cell-scaffold interactions [Baaijens et al., 2010], bioactivity can be enhanced by incorporate biochemical cues to enhance cell recruitment and differentiation, growth factors (e.g., VEGF, HGF) can be embedded in microspheres for sustained release [Lee et al., 2011], scaffolds are modified with RGD peptides or laminin to enhance cell adhesion [Hersel et al., 2003)]. Progenitor cells or MSCs are pre-seeded within the scaffolds to facilitate the process of regeneration [De Coppi et al., 2006].

### Regeneration Patterns in Tendons, Muscles, and Nerves

As analyzed before in this study, the variation pattern of degradation rate and mechanical strength of a biomimetic implant is dependent on the target tissue. Similarly, its regeneration patterns of tendons, muscles, and nerves are distinct.

As depicted in Figure 15, the repairing VML commences at the anastomosis site neighboring autologous muscles. This regeneration process is primarily facilitated by satellite cells (muscle stem cells) situated within the surrounding muscle tissue. These cells are positioned between the outer membrane of the muscle fiber and its basal membrane. This muscle regeneration highlights the inherent regenerative capacity of muscles themselves. Satellite cells absorb and assimilate collagen materials, proliferate, and differentiate into muscle fibers, ultimately repairing the defect.

**Figure 15.**
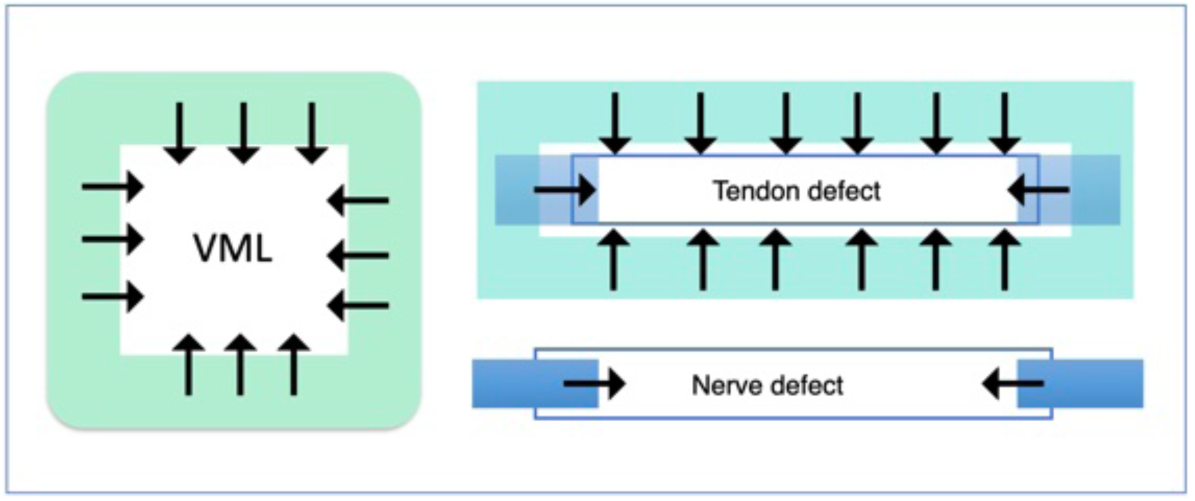
**the diverse regeneration patterns in the repair of VML, tendon defects, and nerve defects. The arrows indicate functional cells moving directions (functionally regenerating direction)**.

In contrast, tendon regeneration originates from paratendon tissue surrounding the defect, rather than the regenerative capacity of the residual autologous tendon. This process is primarily driven by the repair and reconstruction of the surrounding connective tissue, as well as the proliferation activities of capillaries and fibroblasts, and the synthesis of novel collagen fibers, which result in severe adhesion (Figure 15).

Nerve regeneration exhibits a distinct pattern from tendon regeneration and muscle regeneration. Following damage, undamaged neurons serve as the substrate for axon regeneration by augmenting protein synthesis. Axons subsequently progress toward their original target tissue. Simultaneously, Shiwan cells establish a guide channel, facilitating the precise extension and reconnection of axons (Figure 15).

As illustrated in Figure 15, stem cells or progenitor cells are located surrounding or within functional tissues such as tendons, muscles, or nerve tissues. These stem cells possess distinct characteristics, occupy unique locations, and undergo diverse pattern of migration and differentiation, this biomimetic collagen material exhibits same functionality, effectively attracting the polymerization of functional cells to the defect site and facilitating functional tissue regeneration due to its exceptionally low antigenicity and high affinity.

### Future application of the biomimetic materials

This collagen biomimetic material exhibits remarkable regenerative potential in tissue engineering. It will also play a pivotal role in fabricating cell sheets in the field of regenerative medicine of induced pluripotent stem cells (iPS cells) by replacing temperature-sensitive polymer materials (PIPAAm). The technology of PIPAAm has consistently made significant contributions to the formation of cell sheets [Okano et al., 1993]. In Japan, scientific researchers have successfully differentiated iPS cells into cell sheets of retina, cornea, and myocardial membrane [Mandai et al., 2017., Nishida et al., Shimizu et al., 2006]. These cell sheets have been clinically validated and exhibit substantial potential in treating some diseases.

The cultivation of cells on PIPAAm materials facilitates the formation of cell sheets. However, the cooling and detachment of the cell sheet from PIPAAm material can have detrimental effects on cells and complicate the procedures. In contrast, the affinity, nutritional value, and antigenicity of biomimetic collagen materials are significantly superior to those of PIPAAm materials. Substituting PIPAAm with these biomimetic collagen materials presents an optimal carrier for iPS cell growth and differentiation. Moreover, this approach eliminates the need for cooling and detachment, thereby facilitating functional differentiation of cells and tissue regeneration simultaneously improving therapeutic effect.

This advanced material technology based on soluble collagen has demonstrated remarkable efficacy in the functional regeneration. The collagen biomimetic materials can be processed into various colors, shapes and textures with high mechanical strength, including bands, membranes, sponges, strips, and tubes, enabling them to effectively repair a wide range of soft tissue defects (Fig 16).

**Figure 16.**
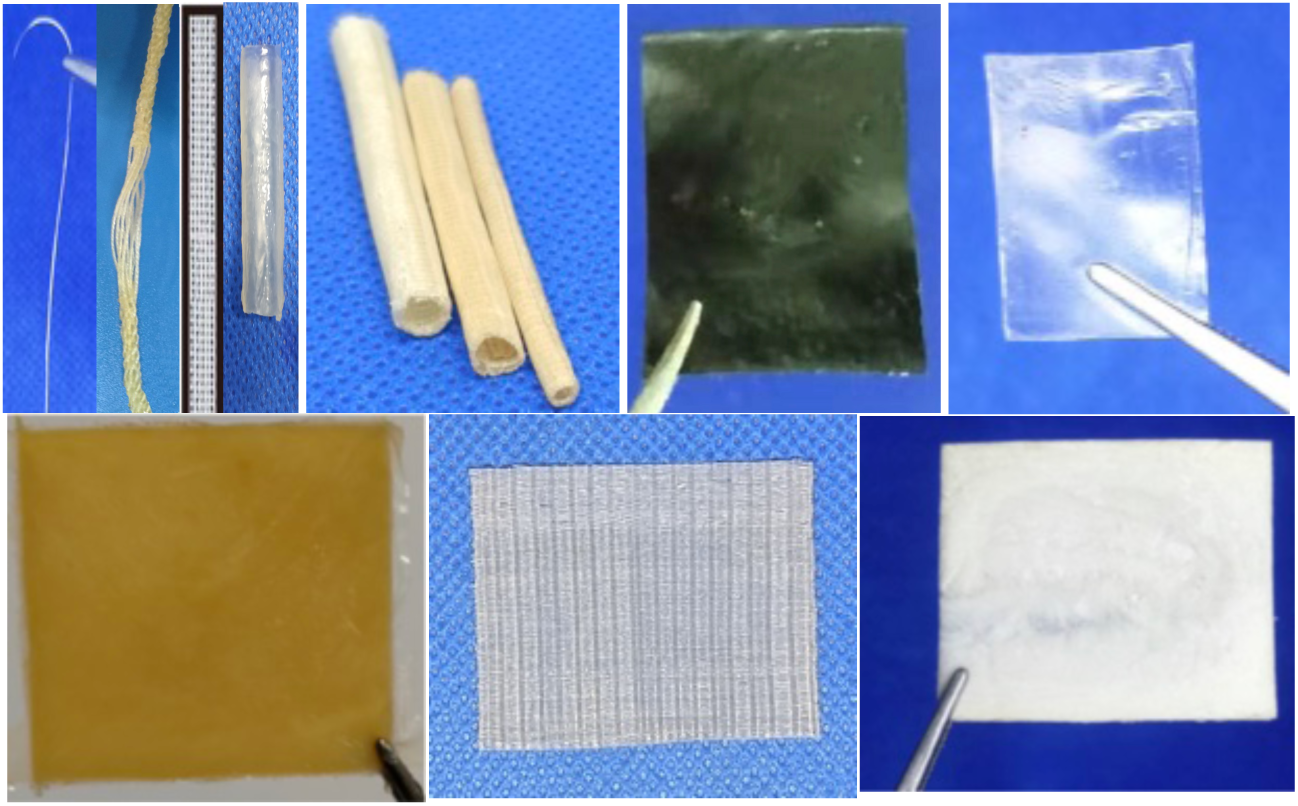
A diverse range of colors, shapes, textures of biomimetic collagen materials.

For instance, the creation of artificial blood vessels and esophagi still presents a significant challenge due to the scarcity of suitable materials. In this study, biomimetic collagen material emerges as the most suitable option for this purpose, as it possesses the substantial potential of regenerating these essential tissues.

The new materials technology also holds substantial medical potential in treating neuromuscular diseases such as, Duchenne muscular dystrophy (DMD), amyotrophic lateral sclerosis (ALS), and spinal muscular atrophy (SMA) by <u>synergizing</u> with advanced technologies, such as CRISPR-Cas9, AAV gene therapy, stem cell technology. One method is to cultivate stem cells (e.g., neural stem cells, neural progenitor cells, or satellite cells) in the biomimetic materials and differentiate them to be muscle fibers or neuromuscular tissue in vitro or in vivo, and subsequently transplanting them into human bodies.

This regenerative scaffold material is not a tissue substitute but a tissue regeneration promoter. It activates and releases biological potential to guide tissue formation and reconstruction, ultimately resulting in the functional regeneration of various tissues.

### Research limitations and future study

Despite the remarkable outcomes, several limitations need to be addressed in this study:(1).Slow Regeneration: Repairing VML, tendon, and nerve defects is relatively slow, necessitating manual intervention to accelerate it. This can be achieved through in vitro implantation of cells (functional cells or stem cells) and growth factors (2).Model Suitability: The present study is confined to the rabbit model. The artificial implants should be further explored for repair in large animals, including sheep, pigs, and humans.(3).Regeneration and Observation Period: The regeneration period is insufficient. An extended observation period is necessary to evaluate the long-term stability of the regenerated tissues. (4).Sports Conditions: In all regeneration cases, rabbits were housed in a small cage. It is imperative to augment their exercise to facilitate and enhance the repair (regeneration) process. (5).Regeneration in Other Body Parts: Muscles and tendons (ligaments) are crucial loading tissues that support the body. Their functional regeneration is equally important in other body parts.(6).Lack of Molecular Mechanisms and Signaling Pathways:The study primarily focuses on histological and biomechanical evaluation, neglecting the crucial role of molecular mechanisms and signaling pathways in understanding the regeneration process. Future research should prioritize the analysis of relevant gene expression and proteome to gain a more comprehensive understanding of the biological characteristics of regenerated tissues.

In regenerative medicine, the primary objective should be to design an effective scaffold using biomimetic materials that closely resemble the target tissue. Advanced technologies such as electrospinning, 3D bioprinting, enable the precise fabrication of biomimetic scaffolds with specific geometric forms, fiber alignment, porosity, and spatial gradients. Stem cells or functional cells can be pre-cultured in scaffolds in vitro under favorable physiological conditions, including mechanical load, oxygen tension, electrical signals, and biochemical gradients.

## Conclusion

This research has successfully developed a high-strength biomimetic scaffold based on soluble collagen. The materials technology demonstrated remarkable efficacy in repairing extensive defects without the necessity of exogenous cells or growth factors. The regenerated tissue exhibited comparable structural and functional characteristics to the autologous tissue. This finding underscores that the primary factor in regeneration lies not in “reproducing tissue structure,” but rather in “activating the potential of stem cells and establishing a favorable microenvironment.”

This study proposes a novel biomimetic materials, thereby opening promising avenues for the field of tissue engineering.

As a “ready-to-use” medical material, it can be produced on a large scale, its low cost and broad clinical adaptability are anticipated to transform into groundbreaking products in the field of regenerative medicine.

